# SMN deficiency disrupts hepatic mitochondrial iron homeostasis and NRF2-dependent redox control in spinal muscular atrophy

**DOI:** 10.64898/2026.01.08.698518

**Authors:** Sofia Vrettou, Stefan Müller, Brunhilde Wirth

## Abstract

Spinal muscular atrophy (SMA), classically defined as a motor neuron disorder caused by deficiency of the survival motor neuron (SMN) protein, is increasingly recognized as a multi-system disease. Among peripheral organs, the liver, essential for metabolic regulation and xenobiotic processing, remains underexplored despite growing evidence of dysfunction. Defining hepatic contributions may be critical for understanding systemic disease progression and optimizing therapeutic strategies.

Here, we performed integrative profiling of liver pathology in SMA by combining unbiased proteomics from wild-type (WT), heterozygous (HET), and SMA mice at the late symptomatic stage (postnatal day 10; P10) with targeted analyses of mitochondrial function, iron metabolism, and redox homeostasis. To resolve temporal dynamics, key pathways were examined at early symptomatic disease (P5), and the reversibility of identified defects was evaluated following SMN-restoring antisense oligonucleotide (ASO) therapy.

We uncover early (P5) activation of the heme biosynthetic pathway, marked by increased ferrochelatase (FECH), preceding overt metabolic disruption. By P10, SMA liver shows a coordinated loss of mitochondrial Complex II integrity, pathological mitochondrial iron accumulation, disruption of the NRF2-KEAP1 antioxidant axis, and heightened susceptibility to ferroptotic redox stress. ASO treatment robustly restored mitochondrial and redox phenotypes, yet FECH remained elevated, indicating sustained heme pathway activation despite SMN rescue. HET mice displayed mild redox abnormalities, revealing a dosage-sensitive hepatic phenotype.

Collectively, these findings delineate a mitochondrial-iron-redox axis uniquely vulnerable in SMA liver, identify ferroptotic sensitivity as an early and therapeutically responsive feature, and highlight persistent heme biosynthesis activation as a target to complement SMN-directed therapies.

## Introduction

Spinal muscular atrophy (SMA) is a severe genetic neuromuscular disease caused by insufficient levels of survival motor neuron (SMN) protein, leading to progressive motor neuron loss and, in severe cases, multi-organ failure [1, 2]. As approximately half of patients develop the most severe form, SMA type I, the advent of SMN-restoring therapies that primarily target the nervous system and markedly prolong survival has revealed peripheral organ pathology as a major and increasingly relevant clinical challenge [3, 4]. SMN deficiency results from homozygous deletion or loss-of-function mutation of *SMN1,* whereas the nearly identical *SMN2* paralog predominantly generates an unstable exon 7-skipped isoform due to a silent nucleotide change [3, 5, 6]. Because *SMN2* provides only limited functional SMN, its copy number strongly influences disease severity [5, 7]. Although motor neuron degeneration drives the clinical presentation [7], SMN is ubiquitously expressed and required for snRNP assembly, pre-mRNA splicing, mRNA trafficking, selenoprotein synthesis, stress granule formation, and cytoskeletal organization [8]. Consistent with these functions, SMN deficiency disrupts axonal mRNA transport and local translation [9–11], and induces metabolic defects with a prominent mitochondria dysregulation [12–16]. Multi-system involvement, including heart, pancreas, adipose tissue, vasculature, liver and kidney, emerges early, often preceding motor neuron loss [17–20], firmly establishing SMA as a systemic disorder with tissue-specific vulnerabilities.

Despite transformative success of approved SMN-targeted therapies (nusinersen, risdiplam, onasemnogene abeparvovec) [21–25], incomplete functional responses remain common [26–28]. Mitochondrial dysfunction and oxidative stress have been reported in SMA [13, 16, 29–33], yet antioxidant-based interventions have shown limited benefit [29–32].

Liver involvement is among the most consistent peripheral phenotypes in SMA. Severe mouse models display small, iron-laden, poorly matured livers [17], while patients and mice exhibit dyslipidemia and steatosis [34]. Hepatocyte intrinsic SMN loss can drive metabolic pathology [35], and selective hepatic SMN restoration improves systemic outcomes and survival in Smn2B/− mice [36]. Because the liver is central to metabolic regulation, iron handling, and drug metabolism, hepatic dysfunction could influence SMA progression and modify responses to SMN-directed therapies [37, 38].

Here, we address these issues using the Taiwanese SMA mouse model, which lacks endogenous murine *Smn* and survives on two copies of the human *SMN2* transgene, recapitulating severe infantile SMA [39]. We performed untargeted quantitative proteomics in symptomatic P10 SMA liver to define global pathway alterations relative to heterozygous (HET) and wild-type (WT) controls. Because HET mice harbor partial SMN reduction, both control genotypes were analyzed to distinguish disease-specific from dosage-sensitive changes. P5 liver was tested to detect early onset of these defects. Early SMN-restoring antisense oligonucleotide (ASO) treatment used to address whether the abnormalities detected at P10 proteomics in SMA were reversible upon early intervention. This integrated strategy provides the first unbiased proteomic profile of SMA liver in the Taiwanese model and defines redox metabolic signatures associated with SMN deficiency and their partial correction after SMN restoration.

## Methods

### Ethics statement

All animal experiments were conducted in accordance with institutional and governmental regulations and approved by the Landesamt für Natur, Umwelt und Verbraucherschutz Nordrhein-Westfalen (LANUV; approval numbers: 81-02.04.2020.A196, 81-02.04.2019.A017, 81-02.04.2019.A138, §4.23.008, §4.22.002). Mice were housed under specific pathogen-free conditions with a 12-h light/dark cycle and ad libitum access to food and water.

### Animals, genotyping, and disease staging

Experiments were performed using the severe Taiwanese mouse model of SMA [39], maintained on a C57BL/6N background [40]. This model lacks endogenous murine *Smn* and carries two copies of the human *SMN2* transgene, resulting in low SMN expression and severe disease phenotype.

Three genotypes were analyzed: wild-type (WT; *Smn^+/+^),* heterozygous carriers (HET; *Smn^+/−^; SMN2^tg/0^*), and SMA (*Smn^−/−^; SMN2^tg/0^*). HET and SMA mice to 50% each were obtained from the same litters [41], while WT controls were age-matched. Genotyping was performed by PCR on tail DNA collected at sacrifice. Primers are listed in Supplementary Table S1. Two postnatal stages were analyzed: postnatal day 5 (P5; early symptomatic) and postnatal day 10 (P10; symptomatic). Tissues were collected, snap-frozen, and stored at −80 °C until analysis. Each experimental group included at least three biological replicates.

### Antisense oligonucleotide treatment

To generate a mild SMN-restored condition, a subset of HET and SMA pups received a single subcutaneous injection of SMN-targeting antisense oligonucleotide (SMN-ASO; 30 µg in saline) at postnatal day 1 (P1), as previously described [42–44]. The ASO targets the SMN2 intronic splicing silencer N1, promoting exon 7 inclusion and partial restoration of full-length SMN protein [45]. Vehicle-treated littermates served as controls. This low-dose regimen produces partial, sub-therapeutic SMN rescue as previously described [40].

### Proteomics analysis

Liver proteomics was performed on P10 mice. Sample preparation, digestion, and cleanup followed the standardized urea-based workflow of the CECAD Proteomics Core Facility. Peptides were analyzed by data-independent acquisition (DIA) LC–MS/MS on an Orbitrap Exploris 480 with FAIMS. Raw files were processed in DIA-NN [46] using a predicted mouse spectral library, with protein-level quantification filtered at 1% FDR. Downstream analyses were performed in Perseus v1.6.15, including log2 transformation, filtering, imputation, principal component analysis, differential expression, and functional enrichment (GO/KEGG). Protein level quantitative data, differential expression analyses, and statistical outputs are provided in Supplementary Tables S2-S7. Proteomics data have been deposited to the ProteomeXchange Consortium via PRIDE (accession number: PXD070887).

### Network and pathway analysis

Significantly altered proteins (FDR < 0.05, S0 = 0.1) from WT vs SMA and HET vs SMA comparisons were analyzed using STRING (v12.0) and visualized in Cytoscape (v3.10). Functional clustering and pathway enrichment were performed using ClueGO/CluePedia with Benjamini–Hochberg correction. Complete listes used as input for network and pathway analyses are provided in Supplementary Tables S2-S7.

### Proteomics data visualization

Proteomics data visualization was performed using Perseus [47], SRplot [48], InstantClue [49], and InteractiVenn [50] for PCA, heatmaps, functional summaries, volcano plots, and overlap analyses. All statistics originated from Perseus analyses.

### Western blotting

Liver tissues were homogenized in RIPA buffer (50 mM Tris-HCl pH 7.4, 150 mM NaCl, 1% NP-40, 0.1% SDS, 0.5% sodium deoxycholate; Thermo Fisher, #89900) containing protease and phosphatase inhibitors (Roche Complete™ EDTA-free, #11836170001; PhosSTOP™, #4906845001). Equal amounts of protein were separated by SDS-PAGE, transferred to PVDF or nitrocellulose membranes, and detected using standard immunoblotting procedures. Signals were visualized by chemiluminescence using SuperSignal West Pico PLUS (Thermo Scientific, #34580). Loading was assessed using Vinculin, HSP90, β-Actin or total protein reverse staining (Thermo Scientific, #24580). Antibodies are listed in Supplementary Table S1.

### Mitochondria isolation, Blue-Native PAGE and in-gel activity assays

Mitochondria were isolated from liver tissue by differential centrifugation as described by Jha et al. [51]. Blue-Native PAGE (BN-PAGE) was used to assess OXPHOS complex assembly, and Clear-Native PAGE (CN-PAGE) was used for in-gel activity assays of complexes I, II, and IV, following established protocols [51].

### Iron quantification (hemic and non-hemic)

Hemic and non-hemic iron levels were measured in mitochondrial and cytosolic fractions using the optimized STAR protocol [52]. Mitochondria were isolated from liver tissue according to the differential centrifugation procedure of Jha et al. [51] and cytosolic fractions were obtained from the post-mitochondrial supernatant by an additional clarification step (7,000 × g, 4 °C). Non-hemic iron was quantified by ferrozine-based colorimetry following acid extraction, while hemic iron was measured fluorometrically using porphyrin detection with hemin as the calibration standard. Values were normalized to protein content.

### Lipid peroxidation

Lipid peroxidation was quantified in liver tissue using a malondialdehyde (MDA) assay kit (Abcam, ab118970), according to the manufacturer’s instructions. Briefly, ∼10-20 mg of tissue was homogenized on ice in the provided lysis buffer supplemented with BHT using a Peqlab homogenizer, followed by centrifugation to remove insoluble material. The MDA-TBA (Malondialdehyde-Thiobarbituric Acid) adduct was generated by incubation with the developer solution at 95 °C for 60 min, cooled on ice, and absorbance measured at 532 nm. MDA standards were used for calibration, and values were expressed as MDA equivalents (nanomoles) normalized to protein content (milligram of protein).

### Glutathione measurements

Reduced and oxidized glutathione levels were determined using a fluorometric GSH/GSSG detection assay kit II (Abcam, ab205811). Tissues were homogenized on ice with the Peqlab homogenizer in the provided assay buffer, followed by deproteinization according to the kit protocol. GSH and total glutathione (GSH + GSSG) were quantified at Ex/Em = 490/520 nm, and GSSG levels calculated as (total glutathione - GSH)/2. The GSH/GSSG ratio was subsequently determined according to the manufacturer’s protocol.

### Statistical analysis

Statistical analyses were performed using GraphPad Prism 10. Data are presented as mean ± SD. Group comparisons were performed using one-way ANOVA with Tukey’s post hoc test unless otherwise stated. P < 0.05 was considered statistically significant.

## Results

### Global proteomic profiling of SMA liver reveals genotype-specific clustering and coordinated mitochondrial-peroxisomal remodeling

To characterize SMN-dependent alterations in the hepatic proteome, we performed untargeted, label-free quantitative proteomic analysis of liver tissue from wild-type (WT), heterozygous (HET), and SMA mice at postnatal day 10 (P10), allowing comparison of SMA liver to both carrier (HET) and fully SMN-intact (WT) controls (Fig. 1A). Proteomic profiling identified a broad and reproducible set of proteins, providing sufficient depth for comparative and functional analyses.

**Figure 1.**
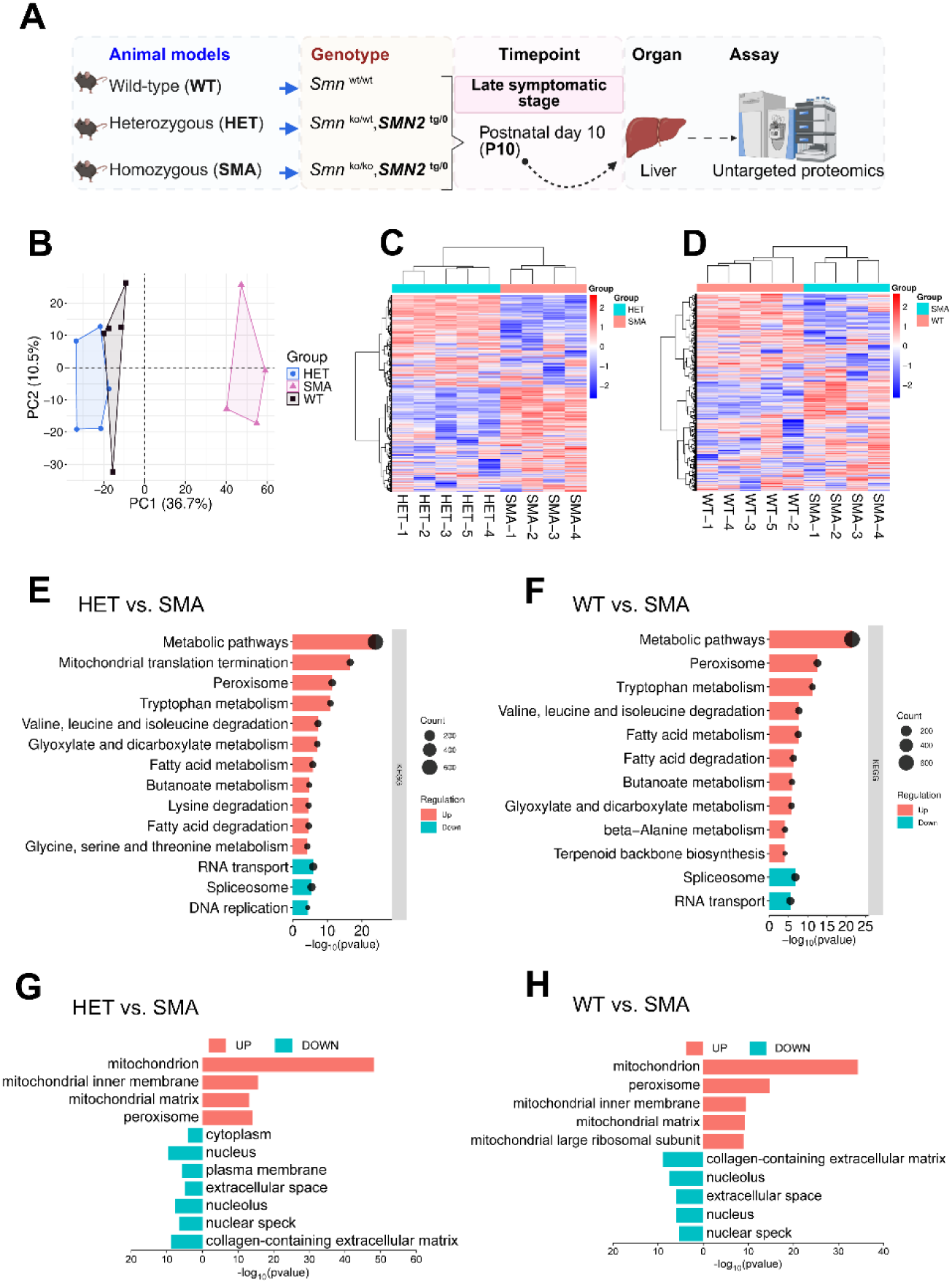
Global proteomic profiling of SMA liver reveals genotype-specific clustering and coordinated organelle remodeling. Liver proteomes from WT, HET, and SMA mice at P10 were analyzed by untargeted label-free proteomics (WT/HET n = 5, SMA n= 4). (A) Experimental overview. (B) PCA showing genotype-driven separation; PC1 and PC2 explain 36.7% and 10.5% of variance, respectively. (C-D) Hierarchical clustering for HET vs SMA and WT vs SMA, grouping primarily by genotype. (E-F) KEGG 1D enrichment analysis of differentially abundant proteins, with pathways enriched among proteins increased in SMA (positive direction) and decreased in SMA (negative direction), selected based on false discovery rate (FDR)- corrected p values and annotation size. (G-H) Gene ontology cellular component (GOCC) enrichment showing predominant mitochondrial and peroxisomal localization of proteins increased in SMA, and enrichment of nuclear, extracellular matrix, and RNA-processing compartments among proteins decreased in SMA, using the same FDR- and size-based selection criteria.

Principal component analysis demonstrated clear genotype-associated segregation of samples, with SMA livers forming a distinct cluster separated from both WT and HET groups (Fig. 1B). In contrast, WT and HET samples exhibited partial overlap, consistent with more modest proteomic differences in heterozygous animals. These relationships were further supported by unsupervised hierarchical clustering, which grouped samples primarily according to genotype and revealed high intra-group consistency (Fig. 1C, D).

Together, these analyses indicate that SMN deficiency is associated with reproducible, genotype-specific remodeling of the liver proteome, establishing a robust framework for subsequent pathway- and network-based investigations.

### Enriched pathways in SMA versus HET or WT

To identify biological processes contributing to the proteomic divergence between genotypes, we performed one-dimensional (1D) enrichment analysis in Perseus using the student’s t-test difference (SMA minus control) for both comparisons (HET vs SMA and WT vs SMA). Complete 1D-enrichment matrices per category of gene ontology annotation are provided in the Data in Brief companion article.

Across both comparisons, KEGG pathway analysis revealed a consistent enrichment of metabolic and organelle-associated programs among proteins increased in SMA (Fig.1E, F). These included broad metabolic pathways, amino acid metabolism, lipid- and fatty acid-related processes, and peroxisome-associated modules, indicating prominent metabolic reprogramming in SMA liver. In contrast, pathways enriched among proteins decreased in SMA were predominantly associated with RNA handling and nuclear processes, including spliceosome- and RNA transport-related categories, with DNA replication emerging selectively in the HET vs SMA comparison.

Gene ontology for cellular component (GOCC) analysis reinforced these patterns (Fig. 1G, H). Proteins increased in SMA were enriched in mitochondrial and peroxisomal compartments across both comparisons, whereas proteins decreased in SMA compared to both HET and WT were predominantly associated with nuclear, extracellular matrix, and RNA-related sub-cellular compartments. Together, these findings indicate that mitochondrial, peroxisomal, and RNA-processing structures account for a substantial proportion of the variance between SMA and control liver proteomes.

To complement the KEGG and GOCC analyses, we also examined molecular function (MF) and Reactome enrichment (Fig. S1). These analyses recapitulated the core trends observed above, with enrichment of metabolic and mitochondrial translation-related terms among proteins increased in SMA and enrichment of RNA binding, mRNA splicing and transcript transport -related categories among proteins decreased in SMA.

Collectively, the reduced representation of spliceosome and RNA-processing categories aligns with established features of SMA [53, 54]. In contrast, the strong, recurrent enrichment of metabolic, mitochondrial, and peroxisomal pathways among SMA-increased proteins in SMA liver highlights an underappreciated component of the SMA hepatic phenotype and motivated the more focused metabolic, mitochondrial, and redox analyses presented in the subsequent sections.

### Differentially abundant proteins in SMA versus HET or WT

Building on the KEGG and GOCC enrichment analyses, which indicated that proteins altered in SMA liver were predominantly associated with metabolic and mitochondrial functions, we next examined the individual protein changes underlying these signatures (Fig. 1E-H). To this end, pairwise differential abundance analyses were performed comparing SMA liver to both HET and WT controls, and results were visualized using volcano plots (Fig. 2A, B).

**Figure 2.**
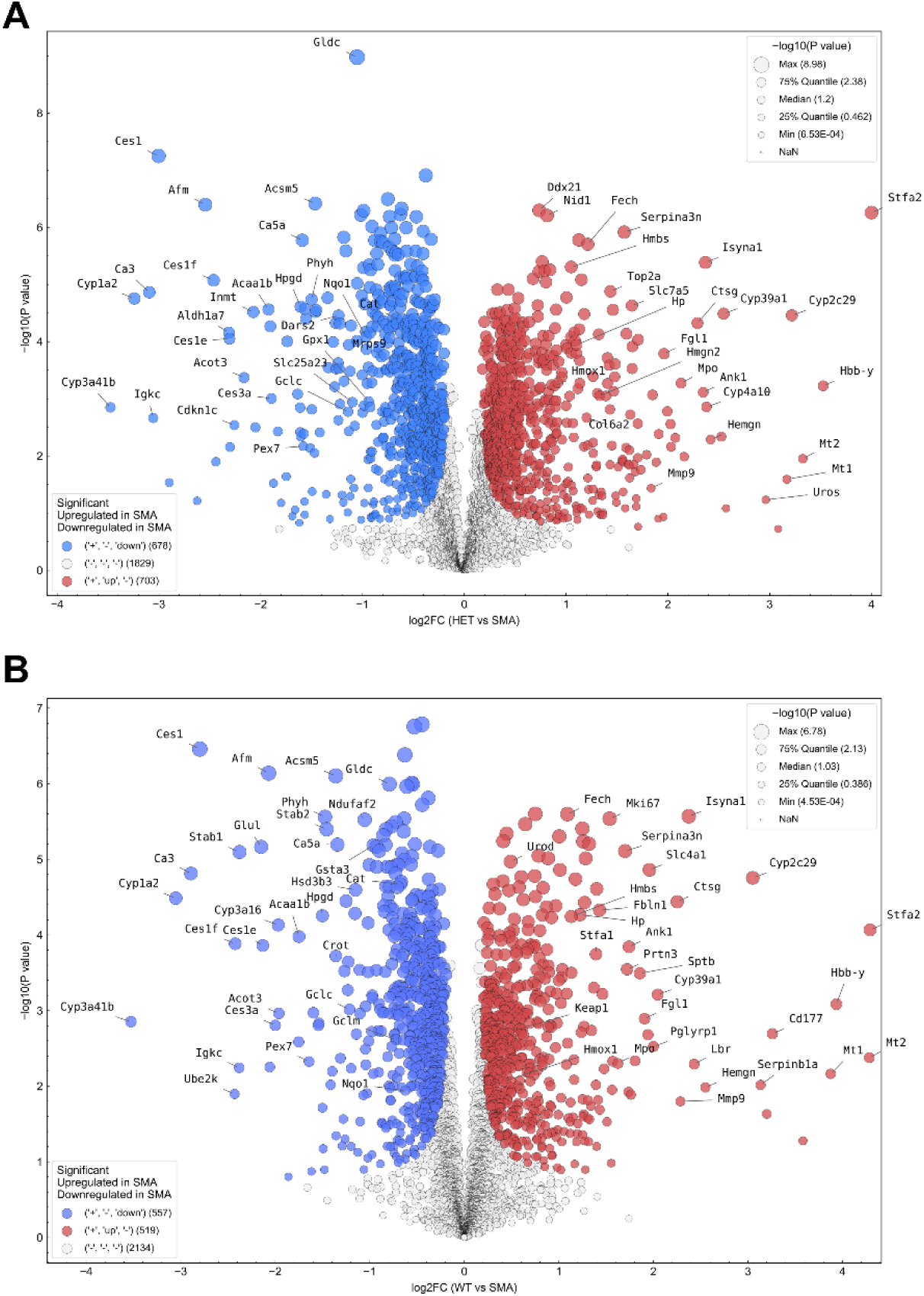
Differential protein abundance in SMA liver compared to HET or WT. (A-B) Volcano plots showing differential abundance between HET vs SMA (A) and WT vs SMA (B) liver proteomes (Student’s t-test, permutation-based FDR < 0.05; S0 = 0.1). Proteins increased in SMA are shown in red and decreased in blue; grey indicates non-significant proteins. Point size reflects –log₁₀(p-value). SMA-elevated proteins were enriched for heme biosynthesis/turnover, redox detoxification, and cytochrome P450 enzymes, whereas SMA-reduced proteins included carboxylesterases and mitochondrial/peroxisomal lipid metabolic enzymes. Many targets were consistently altered across both contrasts, indicating a stable SMA-associated hepatic signature. The corresponding quantitative proteomics data are provided in Supplementary Tables S3-S4.

Both comparisons revealed a consistent pattern of protein abundance changes in SMA liver. Proteins increased in SMA included multiple enzymes involved in heme biosynthesis and turnover, as well as proteins associated with redox and detoxification processes, consistent with a heme- and oxidative stress–related signature. In contrast, proteins reduced in SMA were enriched for carboxylesterases and enzymes involved in peroxisomal and mitochondrial lipid metabolism, indicating impaired hepatic catabolic and peroxisomal capacity.

Importantly, a substantial proportion of significantly altered proteins was shared between the SMA versus HET and SMA versus WT comparisons, with differences largely reflecting the magnitude rather than the direction of change. This consistency suggests that these alterations represent a stable SMA-associated liver signature rather than comparison-specific background variation. Shared and comparison-specific proteins were summarized using Venn diagrams (Fig. S2; Table S2).

Collectively, these analyses indicate that the metabolic and mitochondrial enrichment observed in SMA liver is driven by a coordinated set of proteins linked to heme biosynthesis, heme turnover, and redox-related pathways. The recurrence of these targets across both pairwise comparisons suggested the presence of an underlying coordinated network rather than isolated protein changes in SMA liver, prompting further investigation of their functional relationships.

### Functional enrichment of differentially expressed proteins in SMA liver

To further explore the functional relationships among proteins altered in SMA liver, we extended the analysis from individual protein fold changes to interaction-based network approaches (Fig. 3A). Lists of significantly increased and decreased proteins, together with their corresponding fold-change values derived from the volcano plot analyses, were compiled separately and used as input for STRING to identify known and predicted protein–protein interactions (Fig. 3A).

**Figure 3.**
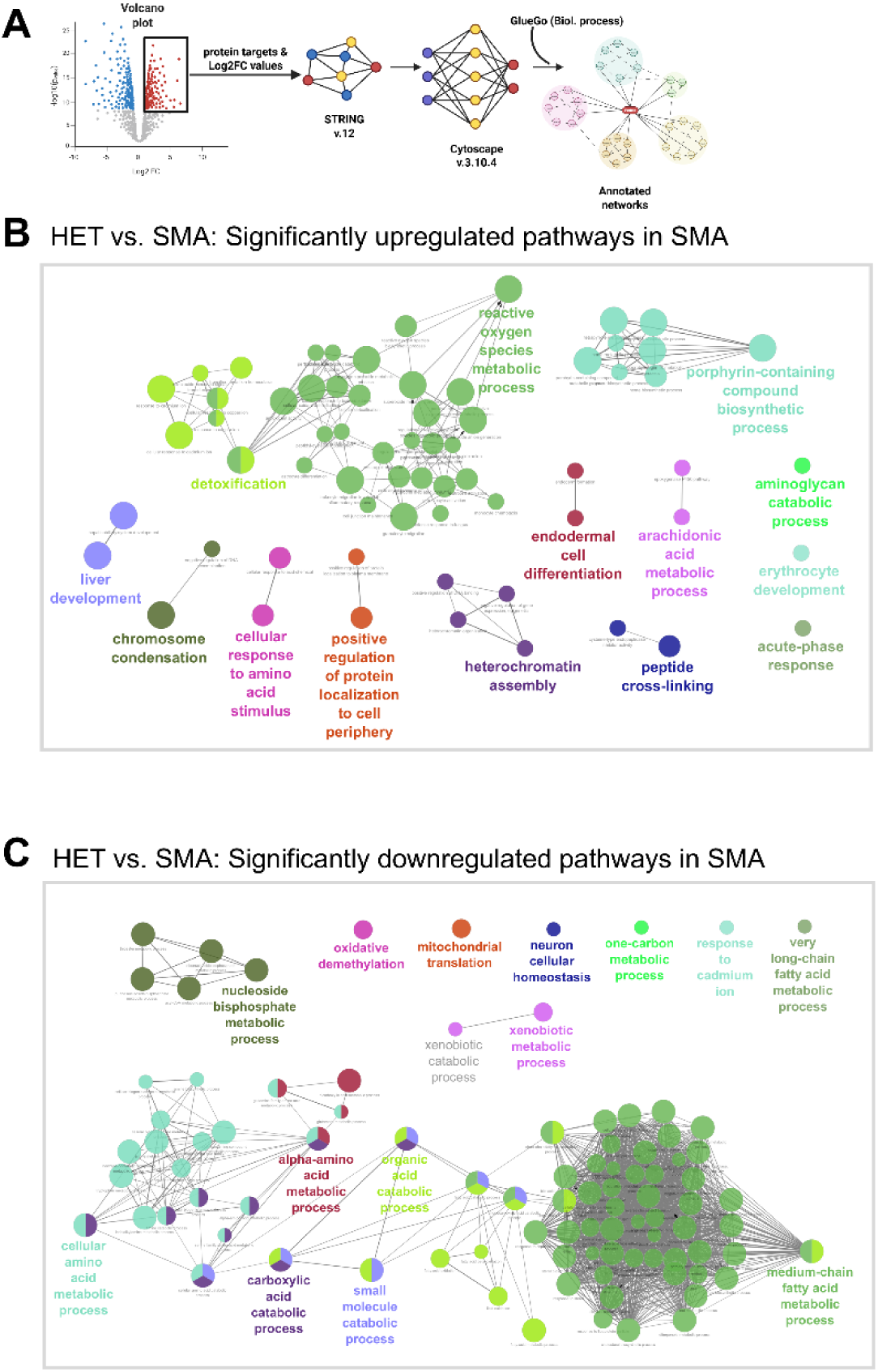
Functional interaction networks highlight SMA-associated redox, metabolic, and porphyrin biosynthesis pathways. (A) Workflow for generating functional networks from significantly altered proteins. Differentially abundant proteins (upregulated or downregulated) from volcano plots were uploaded to STRING to identify interaction networks and imported into Cytoscape for functional mapping using ClueGO (Biological Process). (B-C) ClueGO networks showing significantly enriched biological processes for proteins increased (B) or decreased (C) in SMA relative to HET (FDR < 0.05; kappa ≥ 0.4). Node size reflects the proportion of proteins contributing to each term; colors indicate functional group assignment. Upregulated clusters were enriched for reactive oxygen species metabolism, porphyrin-containing compound biosynthesis, detoxification, and stress-related processes. Downregulated proteins mapped predominantly to amino-acid and fatty-acid catabolism, one-carbon metabolism, and mitochondrial translation. Together, these networks reveal coordinated SMA-associated remodeling of redox, metabolic, and heme-biosynthetic pathways.

The resulting interaction networks were imported into Cytoscape for visualization and functional annotation. Using the ClueGO plugin, proteins were grouped into functionally related clusters based on enriched biological processes and pathways (Fig. 3A). This approach enabled assessment of whether SMA-associated protein changes preferentially localize to specific functional modules rather than occurring as isolated, unrelated events.

### HET versus SMA: Up and downregulated networks

In the comparison between HET and SMA liver proteomes, proteins increased in SMA were organized into functional clusters associated with redox and metabolic stress–related processes, including reactive oxygen species (ROS) metabolism, porphyrin-containing compound biosynthesis, cellular detoxification, and pathways related to synaptic and chromatin organization (Fig. 3A; Fig. S3A). In contrast, proteins decreased in SMA were predominantly enriched in metabolic pathways, including amino acid and fatty acid catabolism, one-carbon metabolism, and mitochondrial translation (Fig. 3B; Fig. S3B).

Among these networks, the most prominent clusters corresponded to ROS metabolism among proteins increased in SMA and medium-chain fatty acid metabolism among proteins decreased in SMA, as reflected by the relative distribution of functional categories shown in the corresponding pie charts (Fig. S3B; Fig. S4B). These patterns indicate that SMA liver exhibits coordinated alterations in redox-associated and metabolic processes when compared with heterozygous controls.

### WT versus SMA: Up and downregulated networks

We next compared SMA liver proteomes with WT controls to assess whether similar functional patterns emerged relative to a fully SMN-intact background (Fig. S4). As observed in the HET comparison, proteins increased in SMA clustered into pathways related to transition metal ion homeostasis, cellular detoxification, and porphyrin-containing compound biosynthesis (Fig.S4A, Fig.S5A).

Conversely, proteins decreased in SMA relative to WT were predominantly associated with lipid and fatty acid metabolism, organic acid catabolism, and broader small-molecule metabolic pathways (Fig.S4B, Fig.S5B). These findings closely parallel those observed in the HET comparison, indicating that the major functional themes distinguishing SMA liver are largely consistent across both control genotypes.

Across both network analyses, porphyrin-containing compound biosynthetic pathway emerged as a recurrently enriched pathway among proteins elevated in SMA liver. This recurring enrichment underscores that SMA liver is not only subjected to widespread redox and metabolic reprogramming but also harbors a specific defect in the enzymatic machinery of heme biosynthesis. To our knowledge, this represents the first proteome-wide evidence implicating coordinated heme metabolic remodeling downstream of SMN deficiency, thereby providing a mechanistic bridge between altered iron handling and impaired mitochondrial function in SMA liver.

### Validation of heme biosynthetic pathway alterations in SMA liver

To validate the alterations in heme biosynthesis suggested by the proteomic and network analyses, we next performed protein-level assessment of key enzymes within this pathway. In the proteomics datasets, multiple enzymes of this pathway were consistently increased in SMA compared to WT or HET, as shown in the volcano plots (Fig.2), and the STRING networks analyses clearly highlighted an upregulation of porphyrin-containing compound biosynthesis (Fig.3A, Fig. S4A). Guided by these findings, we examined the same pathway in P10 using whole-liver RIPA-extracted protein lysates coupled with Western immunoblotting (Fig. 4).

**Figure 4.**
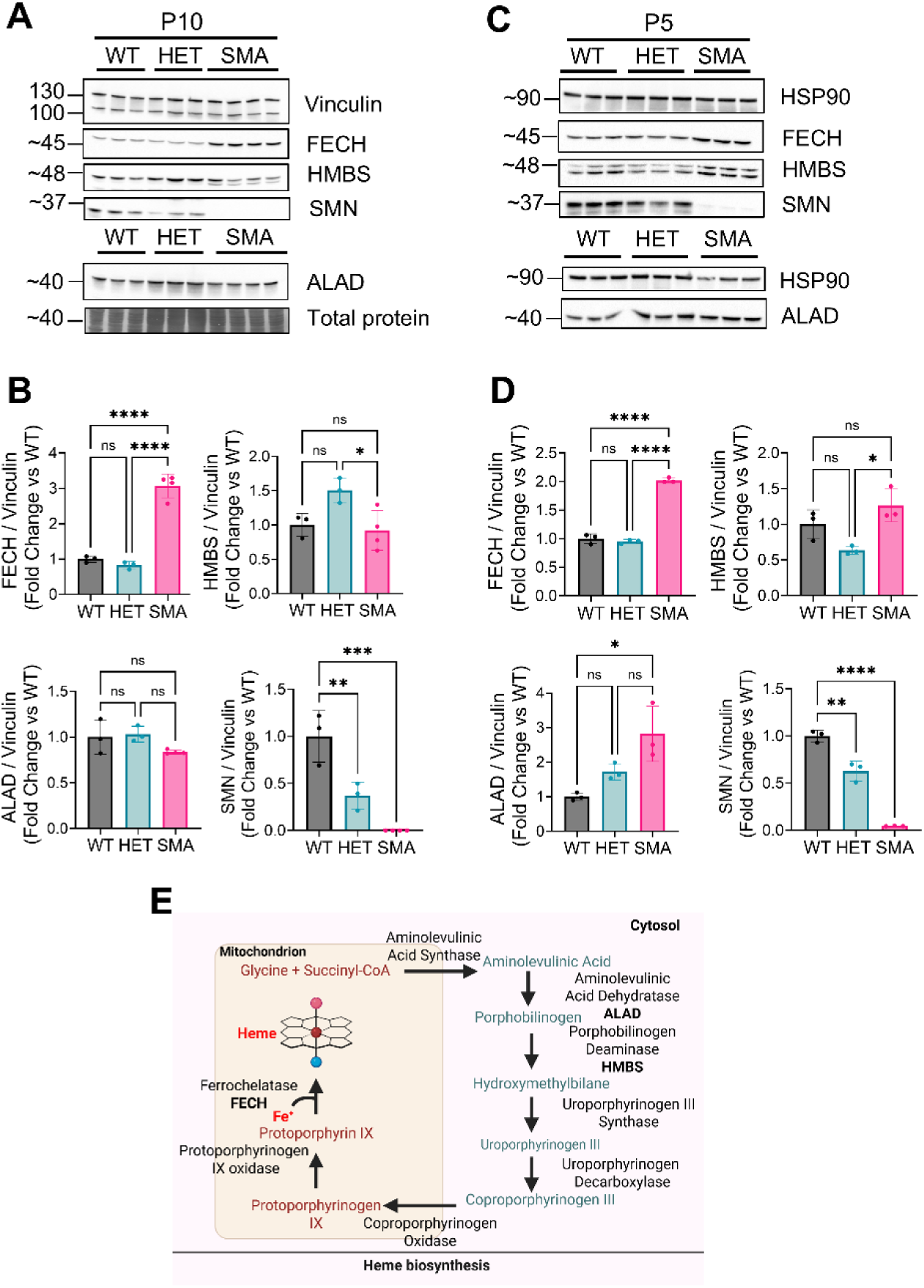
Validation of heme biosynthetic pathway alterations in SMA liver at symptomatic and early symptomatic stages. (A) Representative Western blots of whole-liver lysates from WT, HET, and SMA mice at symptomatic stage postnatal day 10 (P10), probing key heme-biosynthesis enzymes Ferrochelatase (FECH), Hydroxymethylbilane synthase (HMBS), and Aminolevulinate dehydratase (ALAD). Vinculin was used as a loading control. SMN levels are shown as disease reference. (B) Densitometric quantification of FECH, HMBS, ALAD, and SMN protein levels at P10, normalized to Vinculin and expressed relative to WT. FECH is significantly increased in SMA compared to both WT and HET, HMBS is significantly reduced in SMA compared to HET only, and ALAD shows non-significant trends. SMN is markedly reduced in SMA, as expected. (C) Representative Western blots of whole-liver lysates from WT, HET, and SMA mice at the early symptomatic stage postnatal day 5 (P5), probing the same heme-biosynthesis enzymes as in (A). HSP90 was used as a loading control, with SMN included as a disease marker. (D) Densitometric quantification at P5, normalized to HSP90 and compared relative to WT. FECH is already significantly elevated in SMA at this early symptomatic stage. HMBS is increased in SMA relative to HET but not WT, whereas ALAD is increased relative to WT but not HET. SMN shows the expected genotype-dependent reduction in SMA. (E) Schematic overview of the heme-biosynthesis pathway illustrating the cytosolic and mitochondrial steps. Enzymes validated in this study are highlighted, emphasizing that SMA-associated changes are most pronounced at the terminal mitochondrial step catalyzed by FECH. Each dot represents an independent biological replicate. Data are presented as mean ± SD. Statistical analysis was performed using one-way ANOVA followed by Tukey’s multiple-comparisons test. Significance is indicated as p < 0.05 (*), p < 0.01 (**), p < 0.001 (***), and p < 0.0001 (****), ns denotes not significant.

We selected three representative enzymes spanning distinct steps of the cytosolic and mitochondrial arms of the pathway: Ferrochelatase (FECH), Hydroxymethylbilane synthase (HMBS), and Aminolevulinate dehydratase (ALAD) [55]. Notably, FECH showed the strongest statistical significance among heme biosynthesis enzymes in the proteomics dataset, in both HET versus SMA and WT versus SMA comparisons (Fig.2A, B). This increase was confirmed at the protein level, with a marked increase of FECH in SMA versus both WT and HET by immunoblotting at P10 (Fig. 4A, B).

In contrast, cytosolic enzymes upstream of FECH displayed a variable modulation. At P10, HMBS protein levels were reduced in SMA liver compared to HET but not to WT, whereas ALAD levels did not show a significant change (Fig. 4A, B). As an internal disease stage reference, SMN levels were also addressed and showed the expected marked reduction in SMA compared to both WT and HET (Fig. 4A, B). For completeness, individual LFQ profiles for all heme-biosynthesis enzyme detected in the proteome were analyzed (Fig. S6).

Given the pronounced increase of FECH at the symptomatic stage (Fig. 4A, B), we next examined whether this alteration is already detectable at an earlier disease stage. For this, we evaluated the P5 liver, which corresponds to the early symptomatic stage. FECH was already significantly elevated in SMA compared to both WT and HET (Fig. 4C, D), indicating that this mitochondrial step of heme production is increased before the P10 stage. HMBS was significantly higher in SMA compared to HET but not WT, whereas ALAD was significantly elevated in SMA compared to WT but not HET (Fig. 4C, D). SMN expression at P5 showed the anticipated genotype-dependent pattern, with SMA displaying significantly lower levels compared to WT and HET (Fig. 4C, D).

The heme-biosynthesis pathway diagram (Fig. 4E) summarizes the position of these enzymes along the cytosolic–mitochondrial route, illustrating that SMA-associated changes are concentrated at the terminal mitochondrial step catalyzed by FECH. Together, these findings validate the proteomic discovery that SMA liver exhibits selective activation of heme-synthesis machinery, particularly at the ferrochelatase step, reinforcing the idea that alterations in heme production and turnover form a core feature of the symptomatic and early symptomatic stage of the SMA liver metabolic phenotype.

### Coordinated remodeling of hepatic iron handling in SMA liver

Following validation of heme biosynthesis alterations at the protein level, we next examined whether broader iron-associated pathways exhibited coordinated changes in the SMA liver proteome. To this end, iron-related proteins identified in the proteomics dataset were grouped into functional modules encompassing heme-utilizing enzymes, heme-dependent mitochondrial complexes, iron–sulfur cluster biogenesis, and iron acquisition and trafficking (Fig. S7).

Across these modules, SMA liver displayed a consistent pattern indicative of altered iron handling and redox adaptation. Proteins involved in heme catabolism and scavenging including heme oxygenase 1 (HO1), hemopexin (HEMO), haptoglobin (HPT), and biliverdin reductase B (BLVRB) were consistently elevated in SMA liver (Fig. S7A). HO-1 upregulation is especially notable: because heme-biosynthesis enzymes (particularly FECH) are increased in SMA (Fig. 4), the elevation of HO1 indicates that the liver senses an augmented heme burden and actively channels excess heme into biliverdin (via HO1) and bilirubin (via BLVRB) detoxification pathways [56]. The parallel increases in HEMO and HPT further support enhanced scavenging of free heme or heme–protein complexes [57], suggesting higher turnover or influx of heme-derived cargo in SMA. In contrast, components linked to mitochondrial iron utilization and iron– sulfur cluster homeostasis (Fig.S7D, E) exhibited more heterogeneous, yet directional, changes, including iron-sulfur cluster assembly protein 2 (ISCA2) upregulation and ferredoxin reductase (FDXR) downregulation. In parallel, iron uptake and transport pathways were selectively increased (Fig. S7E), including increased expression of transferrin receptor 1 (TFR1).

In summary, these proteomic signatures reveal that SMA liver exhibits a coordinated remodeling of iron handling that extends far beyond heme biosynthesis. The observed pattern is consistent with increased heme turnover, redox stress, and compensatory iron redistribution, providing a systems-level context for the mitochondrial and iron-dependent phenotypes examined in subsequent sections.

### Mitochondria stoichiometry, function and biogenesis in SMA liver

Building on the alterations in heme biosynthesis and the coordinated remodeling of iron and Fe-S cluster-associated pathways, we next examined the impact of these changes on mitochondrial organization, function, and biogenesis in SMA liver. Because both heme and Fe-S clusters serve as essential cofactors for respiratory chain assembly, stability, and activity [58, 59], the proteomic patterns observed suggest that mitochondrial homeostasis may be challenged in the SMA liver. We therefore investigated whether these molecular alterations are associated with changes in oxidative phosphorylation (OXPHOS) complex stoichiometry, respiratory function, and mitochondrial biogenesis.

### Selective impairment of mitochondrial complex II in P10 SMA liver

To assess the status of the OXPHOS system in symptomatic P10 liver, we first examined whole-liver lysates using an OXPHOS antibody cocktail detecting representative subunits of respiratory Complexes I-V (Fig. 5A). At the level of whole-tissue homogenates, a significant reduction of the Complex II subunit SDHB (succinate dehydrogenase complex iron sulfur subunit B) was observed in SMA compared to HET, whereas the abundance of other complexes appeared largely preserved (Fig. 5B).

**Figure 5.**
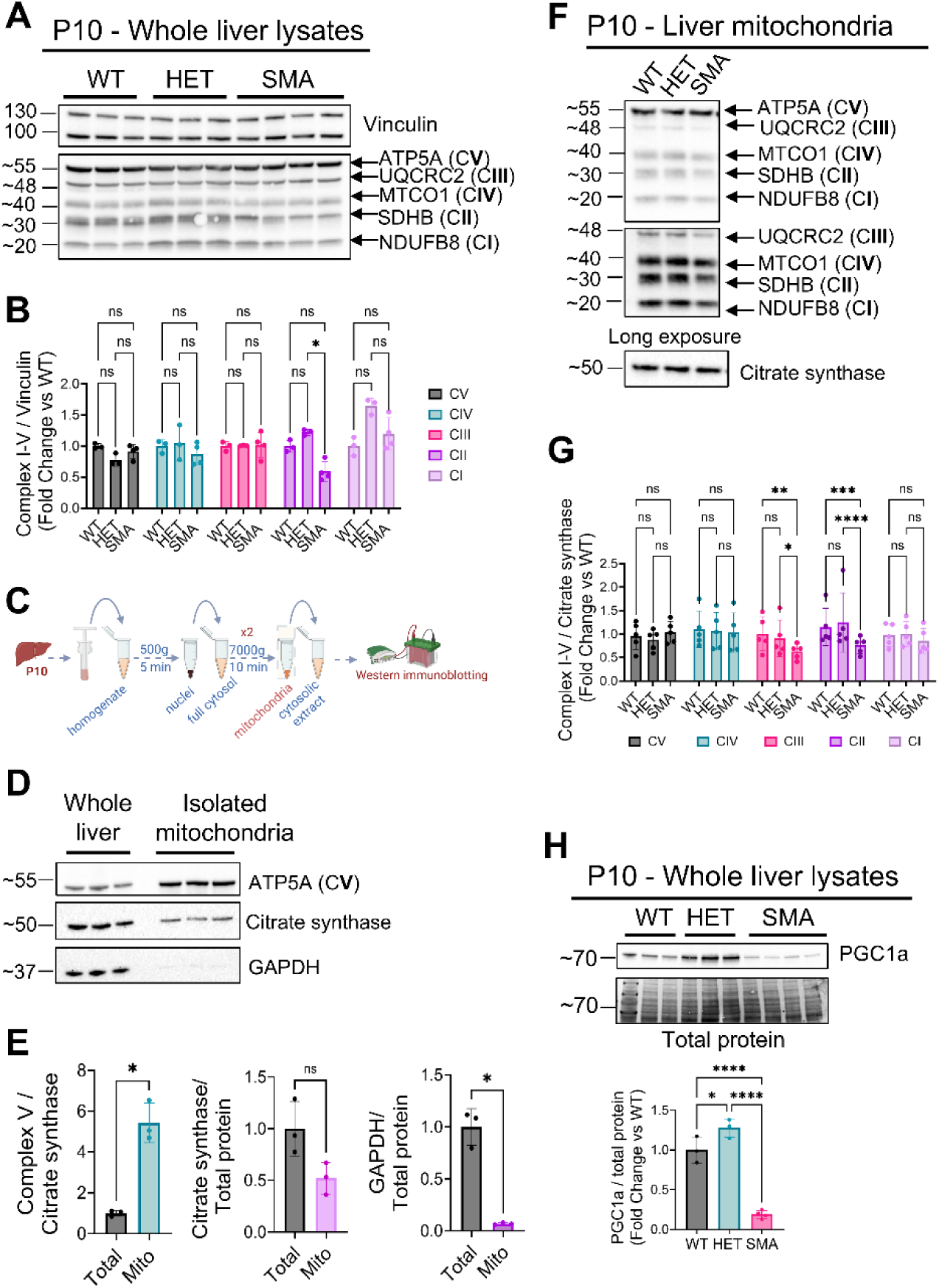
Selective impairment of mitochondrial Complex II and reduced mitochondrial biogenesis in SMA liver. (A) Representative immunoblots of whole-liver homogenates from WT, HET, and SMA mice at postnatal day 10 (P10) probed with an OXPHOS antibody cocktail detecting representative subunits of respiratory Complexes I-V. Vinculin was used as a loading control. (B) Densitometric quantification shows a selective reduction of the Complex II subunit SDHB in SMA, whereas other complexes remain unchanged. (C) Schematic overview of the experimental workflow for liver mitochondrial isolation from P10 mice by differential centrifugation, followed by immunoblot-based analysis. (D) Immunoblot analysis comparing whole-liver lysates and isolated mitochondrial fractions, demonstrating mitochondrial enrichment by increased levels of ATP5A and citrate synthase and depletion of the cytosolic marker GAPDH in mitochondrial preparations. (E) Quantification of mitochondrial enrichment and fraction purity, showing increased Complex V, unchanged citrate synthase levels relative to total protein in isolated mitochondria, and depletion of GAPDH compared to whole-liver lysates. (F) Representative immunoblots of isolated liver mitochondria from WT, HET, SMA, and SMA+ASO mice probed with the OXPHOS antibody cocktail. A long exposure is shown below to enhance visualization of subunits. Citrate synthase was used as a mitochondrial loading control. (G) Densitometric quantification of OXPHOS subunits in isolated mitochondrial fractions normalized to citrate synthase. A robust reduction of the Complex II subunit SDHB is observed in SMA, accompanied by a significant decrease in Complex III subunit UQCRC2, whereas Complexes I, IV, and V remain largely unchanged. (H) Representative immunoblots and quantification of PGC1α protein levels in P10 liver normalized to total protein. PGC1α is significantly reduced in SMA compared to WT and HET, whereas HET shows increased PGC1α relative to WT. Each dot represents an independent biological replicate. Data are presented as mean ± SD. Statistical analysis was performed using one-way ANOVA followed by Tukey’s multiple-comparisons test. Significance is indicated as p < 0.05 (*), p < 0.01 (**), p < 0.001 (***), and p < 0.0001 (****), ns denotes not significant.

To determine whether this defect was more pronounced at the level of mitochondria, liver mitochondria were isolated from P10 animals using differential centrifugation according to the Jha et al protocol [51] (Fig. 5C). Subfractionation efficiency and mitochondrial enrichment were verified by immunoblotting for mitochondrial markers, including ATP5A (Complex V) and citrate synthase, together with depletion of the cytosolic protein GAPDH relative to whole-liver lysates (Fig. 5D, E), confirming successful isolation of mitochondrial subfractions.

Immunoblotting of isolated mitochondrial fractions with the OXPHOS antibody cocktail revealed a robust reduction of the Complex II subunit SDHB in SMA compared to WT and HET (Fig. 5F, G). In addition, mitochondrial enrichment uncovered a significant decrease in the Complex III subunit UQCRC2 in SMA (Fig. 5F, G), a difference that was not readily detected in whole-liver lysates. In contrast, the abundance of subunits representing Complexes I, IV, and V remained largely unchanged across groups. These findings indicate that mitochondrial isolation enhances detection sensitivity and reveals selective vulnerabilities within the respiratory chain that are partially masked in whole-tissue homogenates.

Collectively, these data identify Complex II as the most consistently and strongly affected respiratory complex in P10 SMA liver, with additional involvement of Complex III emerging upon mitochondrial enrichment. Given the central role of Complex II at the interface of the tricarboxylic acid cycle and the electron transport chain [60], and its dependence on iron–sulfur cluster and heme cofactors [61], this selective impairment further supports a link between mitochondrial dysfunction and altered redox–iron metabolism in SMA liver.

### PGC1α is downregulated in P5 and P10 SMA liver

Because mitochondrial biogenesis is a key determinant of respiratory chain content [62], we next assessed hepatic PGC1α abundance. PGC1α levels were significantly reduced in SMA liver at P10 relative to both WT and HET controls (Fig. 5H). In contrast, P10 HET liver exhibited a significant increase in PGC1α abundance compared to WT (Fig. 5H), indicating differential regulation of mitochondrial biogenesis programs in the context of partial versus severe SMN deficiency.

To further examine the temporal relationship between mitochondrial biogenesis and respiratory chain composition, the same analyses were performed at the earlier P5 timepoint. At this stage, PGC1α abundance was already reduced in SMA liver, whereas the steady-state levels of individual OXPHOS complexes remained unchanged, as assessed by immunoblotting (Fig. S8 A, B).

The absence of overt respiratory subunit loss at P5 indicates that alterations in mitochondrial biogenesis precede structural degeneration of the electron transport chain, placing PGC1α downregulation upstream in the trajectory leading to the selective Complex II impairment observed at the symptomatic stage.

### Progressive dysregulation of the NRF2-KEAP1 antioxidant axis in SMA liver

In light of the evidence for redox imbalance, altered iron and heme flux, and impaired mitochondrial function in SMA liver, we next examined the status of the NRF2-KEAP1 antioxidant axis, a central transcriptional regulator of cytoprotective responses [63, 64]. At the symptomatic P10 stage, SMA liver exhibited a marked reduction of NRF2 protein accompanied by increased KEAP1 abundance, consistent with a state of NRF2 repression (Fig. 6A-C). This was paralleled by decreased expression of canonical NRF2 targets [65], including NQO1, GCLC, and the lipid-peroxide detoxifying enzyme GPX4 (Fig. 6D-G), indicating a collapse of NRF2-driven antioxidant and glutathione-dependent defense pathways.

**Figure 6.**
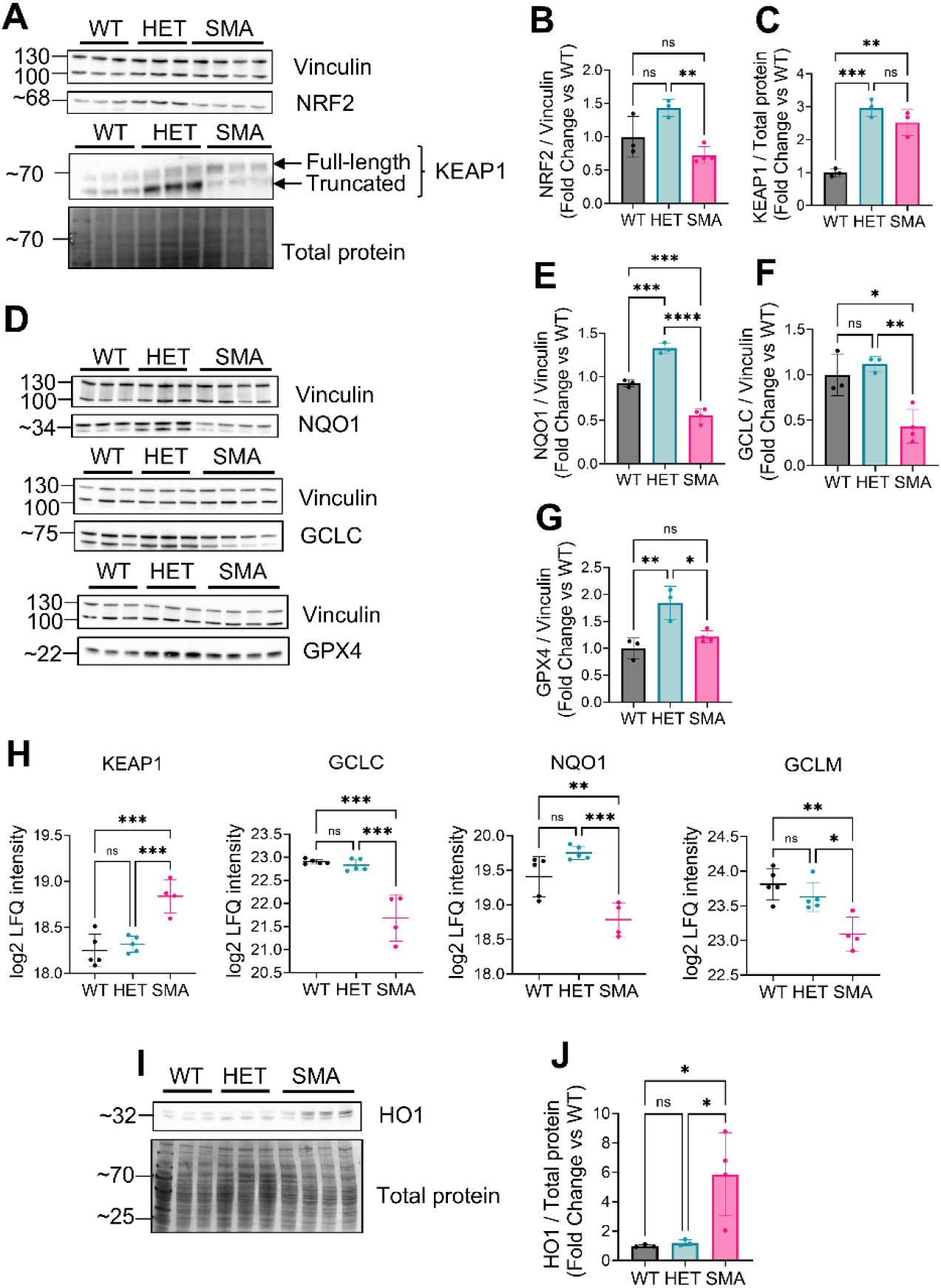
Dysregulation of the NRF2-KEAP1 antioxidant axis in symptomatic SMA liver. (A) Representative Western blots of whole-liver lysates from WT, HET, and SMA mice at postnatal day 10 (P10) showing protein levels of NRF2 and KEAP1. Vinculin or total protein staining was used as loading control, as indicated. (B-C) Densitometric quantification of NRF2 and KEAP1 protein levels at P10, normalized to the corresponding loading control and expressed relative to WT. SMA liver shows reduced NRF2 and increased KEAP1 abundance, consistent with repression of NRF2 signaling. (D) Representative Western blots of the canonical NRF2 target proteins NQO1, GCLC, and GPX4 in WT, HET, and SMA P10 liver lysates. Vinculin was used as a loading control. (E-G) Quantification of NQO1, GCLC, and GPX4 protein levels at P10, normalized to Vinculin and expressed relative to WT. All NRF2 targets are significantly reduced in SMA, indicating impaired antioxidant and glutathione biosynthetic capacity. (H) Log₂ LFQ intensities derived from liver proteomics for selected NRF2 axis components (GCLC, GCLM, NQO1, and KEAP1) at P10. LFQ values are shown for proteins that were robustly quantified and reached statistical significance in the proteomics dataset and display the same directionality of change as observed by immunoblotting, thereby corroborating repression of NRF2 signaling at the system level. (I-J) Representative Western blot and corresponding quantification of HO1 protein levels in WT, HET, and SMA P10 liver. HO1 was normalized to total protein. HO1 is markedly increased, showing impaired regulation of lipid-peroxide detoxification and increased heme turnover. Each dot represents an independent biological replicate. Data are presented as mean ± SD. Statistical analysis was performed using one-way ANOVA followed by Tukey’s multiple-comparisons test. Significance is indicated as p < 0.05 (*), p < 0.01 (**), p < 0.001 (***), and p < 0.0001 (****), ns denotes not significant.

These immunoblot findings were supported by corresponding proteomics data. Log₂ LFQ intensities for NRF2 axis components detected in the liver proteome (GCLC, GCLM, NQO1, KEAP1) showed the same directionality of change at P10, further validating the proteomic dataset and corroborating repression of NRF2 signaling at the protein level (Fig. 7H). In contrast, heme oxygenase 1 (HO1), whose expression can be regulated by multiple stress-responsive pathways independent of NRF2 [56], was strongly upregulated (Fig. 6I, J), aligning with the elevated heme turnover detected in the proteomics dataset (Fig. S7A).

**Figure 7.**
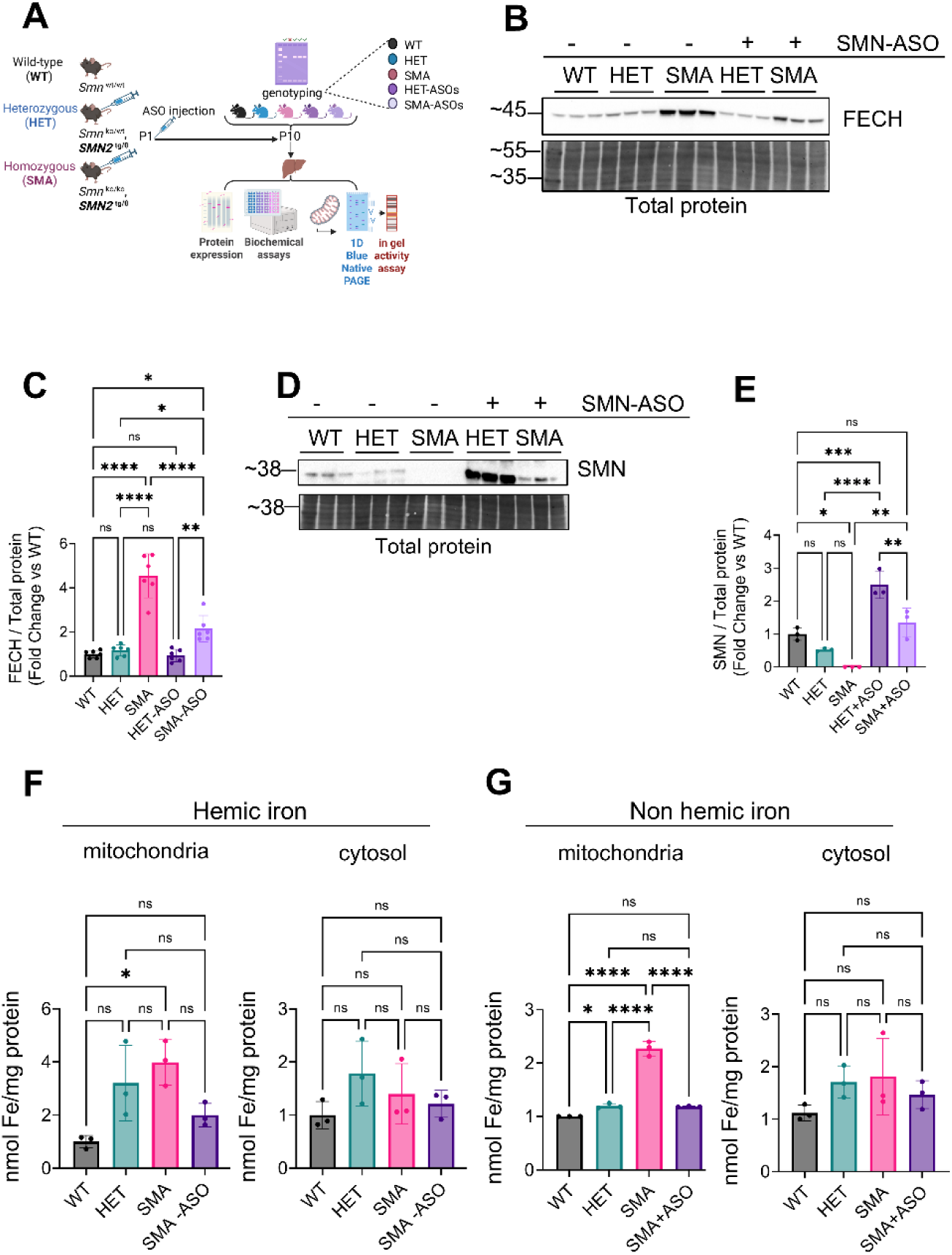
Partial restoration of SMN levels mitigates FECH abundance and mitochondrial iron accumulation in SMA liver. (A) Experimental design schematic. HET, and SMA pups were injected at postnatal day 1 (P1) with a low dose (30µg) of SMN-targeting antisense oligonucleotide (SMN-ASO) or saline, generating five experimental groups (WT, HET, SMA, HET+ASO, SMA+ASO). Liver tissues were collected at P10 for protein expression analyses, biochemical assays, and mitochondrial functional studies. (B-C) Representative Western blot and densitometric quantification of Ferrochelatase (FECH) protein levels in WT, HET, and SMA liver with or without SMN-ASO treatment at P10. FECH abundance is significantly reduced in SMA+ASO compared to untreated SMA, while remaining elevated relative to WT and untreated HET. FECH levels in HET are not altered by ASO administration. SMN protein levels are increased in both SMA and HET following ASO treatment. Total protein staining was used as a loading control. (D-E) Representative Western blot and densitometric quantification of SMN protein levels in WT, HET, and SMA liver with or without SMN-ASO treatment at P10. SMN protein levels are increased in both SMA and HET following ASO treatment. Total protein staining was used as a loading control. (F) Hemic iron levels were biochemically measured in isolated mitochondrial and cytosolic fractions from WT, HET, SMA, and SMA+ASO livers at P10. SMA shows increased mitochondrial hemic iron with no change in the cytosolic fraction, which is fully normalized by SMN-ASO treatment. (G) Non-hemic iron levels were biochemically measured in isolated mitochondrial and cytosolic fractions from the same groups. SMA exhibits a marked increase in mitochondrial non-hemic iron, with cytosolic levels remaining unchanged. SMN-ASO treatment prevents mitochondrial iron accumulation and restores iron levels to those observed in WT and HET animals. Each dot represents an independent biological replicate. Data are presented as mean ± SD. Statistical analysis was performed using one-way ANOVA followed by Tukey’s multiple-comparisons test. Significance is indicated as p < 0.05 (*), p < 0.01 (**), p < 0.001 (***), and p < 0.0001 (****); ns denotes not significant.

In line with impaired NRF2 signaling and reduced GPX4 abundance, a network-level analysis of ferroptosis-associated proteins detected in the liver proteome revealed a coordinated involvement of glutathione metabolism, iron uptake and transport, and mitochondrial-associated processes (Fig. S9).

To determine whether these abnormalities arise early, we assessed the P5 stage and found that NRF2 and KEAP1 levels were still unchanged, whereas NQO1 and GCLC were already elevated (Fig. S10), consistent with intact or compensatory activation of NRF2 signaling. Notably, GPX4 was significantly reduced in SMA even at P5 (Fig. S10), indicating an early deficit in lipid-peroxide detoxification that precedes the later repression of NRF2 signaling. On the other hand, HO1 expression at P5 was unchanged across groups (Fig. S10), further supporting the view that heme-related stress responses emerge only later in the disease course.

Together, these findings reveal a temporal progression in which SMA liver initially engages NRF2-dependent antioxidant defenses alongside an early reduction in GPX4, before transitioning at later symptomatic stages into repression of the NRF2-KEAP1 axis.

### SMN restoration partially normalizes FECH levels and mitochondrial iron accumulation in SMA liver

To determine whether the heme and iron related abnormalities identified in SMA liver are responsive to SMN restoration, we next employed a mild SMN-restoring intervention. SMA and HET pups were injected at P1 with a low dose of an SMN targeting antisense oligonucleotide (SMN-ASO), generating five experimental groups (WT, HET, SMA, HET plus ASO, SMA plus ASO) that were assessed at P10 for protein level, biochemical measurements, and mitochondrial parameters (Fig. 7A). This experimental design allowed us to evaluate whether partial restoration of SMN levels, comparable to levels achieved in SMA type 1 patients treated with SMN-enhancing therapies, is sufficient to modulate the elevated heme synthesis and iron handling defects observed in untreated SMA livers.

We first examined the terminal mitochondrial enzyme of heme biosynthesis, ferrochelatase (FECH) (Fig. 7B, C), which was previously shown to be markedly increased in SMA liver (Figs. 2 and 4). SMN-ASO treatment was associated with significant reduction in FECH abundance in SMA mice relative to untreated SMA animals, with FECH levels returning to the range observed in HET controls, although remaining elevated compared to WT (Fig. 7B, C). FECH levels in HET mice were not significantly altered by ASO administration (Fig. 7B, C), indicating that the partial normalization of FECH is selective to the SMA context rather than the general consequence of ASO treatment. This selective response occurred despite SMN-ASO-mediated increases in SMN protein levels in both SMA and HET animals (Fig. 7D, E), suggesting that correction of FECH upregulation is contingent on the pathological state present in SMA liver.

Consistent with the partial normalization of FECH, analysis of the heme stress-responsive enzyme HO1 revealed a reduction in SMA+ASO liver relative to untreated SMA, while HO1 levels remained elevated compared to HET controls (Supplementary Fig. S11). This pattern indicates that SMN restoration attenuates, but does not fully resolve, the heme stress response associated with SMA liver pathology.

Given that increased FECH abundance would be expected to enhance mitochondrial heme synthesis, we next quantified hemic and non-hemic iron in isolated mitochondrial and cytosolic fractions (Fig. 7F, G). SMA liver displayed a significant increase in mitochondrial hemic iron content, whereas cytosolic hemic iron levels were unchanged (Fig. 7F). A similar compartment-specific pattern was observed for non-hemic iron, which showed a strong elevation specifically in SMA mitochondria but not in the cytosolic fraction (Fig. 7G). In addition, HET mitochondria displayed a modest but significant increase in non-hemic iron levels relative to WT (Fig. 7G). Together, these findings indicate that iron accumulation in SMA liver is preferentially localized to mitochondria and affects both heme-incorporated and labile non-heme pools.

Importantly, SMN-ASO treatment was associated with prevention of mitochondrial iron accumulation in SMA mice, restoring both hemic (Fig. 7F) and non-hemic (Fig. 7G) iron levels to those observed in untreated WT and HET controls. These effects occurred despite only partial normalization of FECH abundance, suggesting that mitochondrial iron accumulation and FECH upregulation are related but not strictly coupled processes.

The presence of increased hepatic iron in SMA is consistent with prior reports describing iron dysregulation in SMN-deficient mouse models [17, 66, 67]. However, previous studies primarily relied on histological staining or whole-tissue iron measurements and did not resolve subcellular iron distribution. The current data extend these observations by indicating that iron accumulation in SMA liver is selectively concentrated within the mitochondrial compartment rather than uniformly distributed throughout the cytosol. This compartmentalization provides additional mechanistic context linking SMN deficiency to mitochondrial iron loading, with potential implications for heme biosynthesis, iron-sulfur cluster metabolism, and redox balance.

Together, these results demonstrate that mitochondrial iron accumulation is a key feature of the SMA hepatic phenotype and that partial restoration of SMN expression is associated with normalization of mitochondrial iron levels, while correction of elevated FECH abundance remains incomplete at this stage of disease.

### SMN-ASO selectively restores Complex II integrity and activity but fails to rescue early PGC1α repression in SMA liver

To determine whether SMN restoration is associated with correction of the mitochondrial abnormalities observed in SMA liver, we assessed respiratory chain organization and function in isolated liver mitochondria from WT, HET, SMA, and SMA+ASO treated animals at P10. BNPAGE coupled with OXPHOS immunoblotting, and in gel activity assays were used to evaluate the integrity and enzymatic activity of mitochondrial respiratory complexes (Fig. 8A).

**Figure 8.**
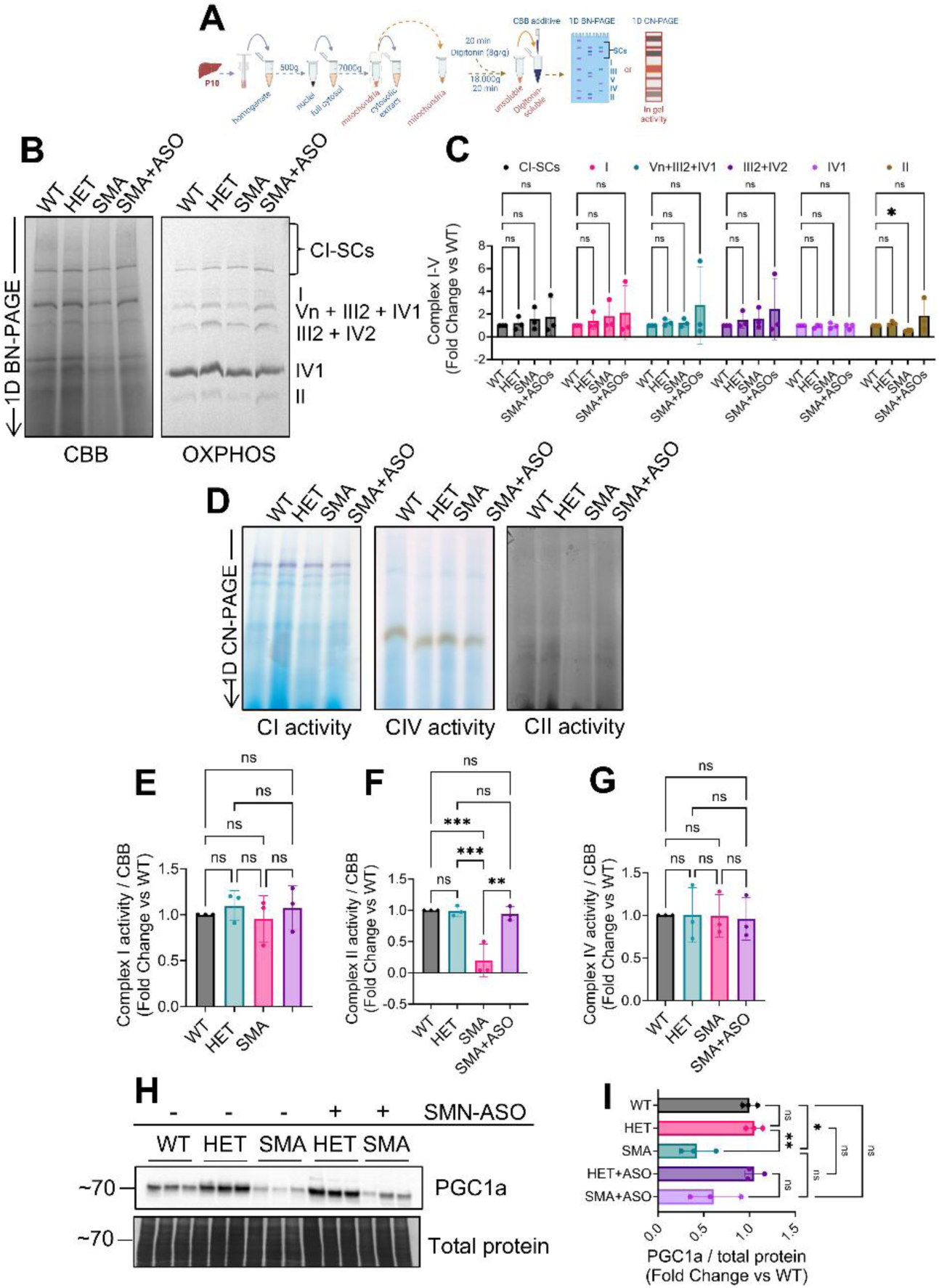
SMN-ASO selectively restores mitochondrial Complex II integrity and activity but fails to rescue PGC1α repression in SMA liver. (A) Schematic overview of the experimental workflow for isolation of liver mitochondria from WT, HET, SMA, and SMN-ASO treated mice at postnatal day 10 (P10), followed by analysis using Blue Native PAGE (BN-PAGE) and Clear Native PAGE (CN-PAGE). (B) BN-PAGE analysis of digitonin-solubilized liver mitochondria visualized by Coomassie Brilliant Blue (CBB) staining and OXPHOS immunoblotting. SMA mitochondria display reduced abundance of Complex II–containing assemblies, whereas SMNASO treatment restores Complex II band intensity to levels comparable to WT and HET animals. The organization of other respiratory complexes and supercomplexes appears largely preserved across groups. (C) Quantification of BN-PAGE–resolved respiratory complexes and supercomplexes expressed as fold change relative to WT. For BN-PAGE OXPHOS immunoblots, the abundance of Complex I, Complex II, and supercomplexes was normalized to Complex IV, which was stable across conditions, whereas Complex IV was normalized to total protein visualized by CBB staining. (D) CN-PAGE in-gel activity assays for Complex I (CI), Complex IV (CIV), and Complex II (CII). Complex II activity is selectively reduced in SMA mitochondria and restored by SMN-ASO treatment, while CI and CIV activities remain comparable across genotypes. (E–G) Quantification of in-gel enzymatic activities for CI (E), CII (F), and CIV (G). For CN-PAGE analyses, CI, CII, and CIV activities were normalized to the CBB signal of Complex I, which served as a stable loading reference across samples. SMN-ASO treatment fully normalizes Complex II activity in SMA mitochondria, whereas CI and CIV activities remain unchanged across all groups. (H) Representative immunoblot of PGC1α protein levels in WT, HET, SMA, HET+ASO, and SMA+ASO liver samples at P10. Total protein staining was used as a loading control. (I) Densitometric quantification of PGC1α normalized to total protein and expressed as fold change relative to WT. PGC1α levels are significantly reduced in SMA liver and are not restored by SMN-ASO treatment, indicating persistent repression of mitochondrial biogenesis despite rescue of Complex II integrity and activity. Each dot represents an independent biological replicate. Data are presented as mean ± SD. Statistical analysis was performed using one-way ANOVA followed by Tukey’s multiple-comparisons test. Significance is indicated as p < 0.05 (*), p < 0.01 (**), p < 0.001 (***), and p < 0.0001 (****); ns denotes not significant.

Consistent with earlier observations (Fig. 5), mitochondria from SMA liver displayed a pronounced reduction in Complex II integrity (Fig. 8B, C), as evidenced by diminished SDHB-containing assemblies. SMN-ASO treatment was associated with restoration of Complex II band intensity in SMA+ASO animals to levels comparable to those observed in untreated WT and HET controls (Fig. 8B, C), indicating recovery of Complex II structural integrity. In contrast, the abundance and organization of other individual complexes and respiratory supercomplexes did not differ appreciably among groups (Fig. 8B, C), supporting the selective vulnerability of Complex II in SMA liver. Functional analyses were consistent with structural findings: In-gel activity assays revealed a significant reduction in Complex II enzymatic activity in SMA mitochondria, which was restored to control levels following SMN-ASO treatment (Fig. 8D, F). Activities of the remaining respiratory complexes were comparable across all experimental groups (Fig. 8D, E, G), in line with their preserved structural integrity and prior observations (Fig. 5).

Given prior evidence suggesting that defects in mitochondrial biogenesis may precede respiratory chain alterations in SMA liver (Fig. 5E, S8B), we next examined the expression of mitochondrial biogenesis regulator PGC1α (Fig. 8H). PGC1α was markedly reduced in SMA liver at P10 (Fig. 8H, I), consistent with our earlier findings at P5 (Fig. S8B). Notably, SMN-ASO treatment did not restore PGC1α levels in SMA+ASO animals (Fig. 8H, I). This persistence of PGC1α repression despite recovery of Complex II integrity and activity suggests that early alterations in mitochondrial biogenesis programs may not be readily reversible by partial SMN restoration at this stage.

Collectively, these data indicate that SMN-ASO treatment is associated with selective recovery of Complex II structure and function in SMA liver, while early repression of PGC1α expression appears to persist, potentially reflecting a distinct or earlier-established component of mitochondrial dysregulation in this tissue.

### SMN-ASO restores NRF2 antioxidant signaling and hepatic redox homeostasis in SMA liver

To determine whether SMN restoration is associated with correction of the repression of the NRF2 antioxidant pathway observed in P10 SMA liver (Fig. 6), we examined key components of the NRF2-KEAP1 axis and downstream antioxidant responses in P10 animals following SMN-ASO administration (Fig. 9A-C). Consistent with earlier findings (Fig. 6), untreated SMA animals showed reduced NRF2 accompanied by increased KEAP1 abundance (Fig. 9A-C). SMN-ASO treatment was associated with a significant increase in NRF2 levels in SMA+ASO mice and a reduction in KEAP1 towards control levels, whereas HET+ASO animals did not show significant changes relative to untreated HET mice (Fig. 9A). These findings indicate that the NRF2-KEAP1 imbalance observed in SMA liver is responsive to SMN restoration and suggest that reduced KEAP1 abundance may contribute to enhanced NRF2 stability in the SMA+ASO context.

**Figure 9.**
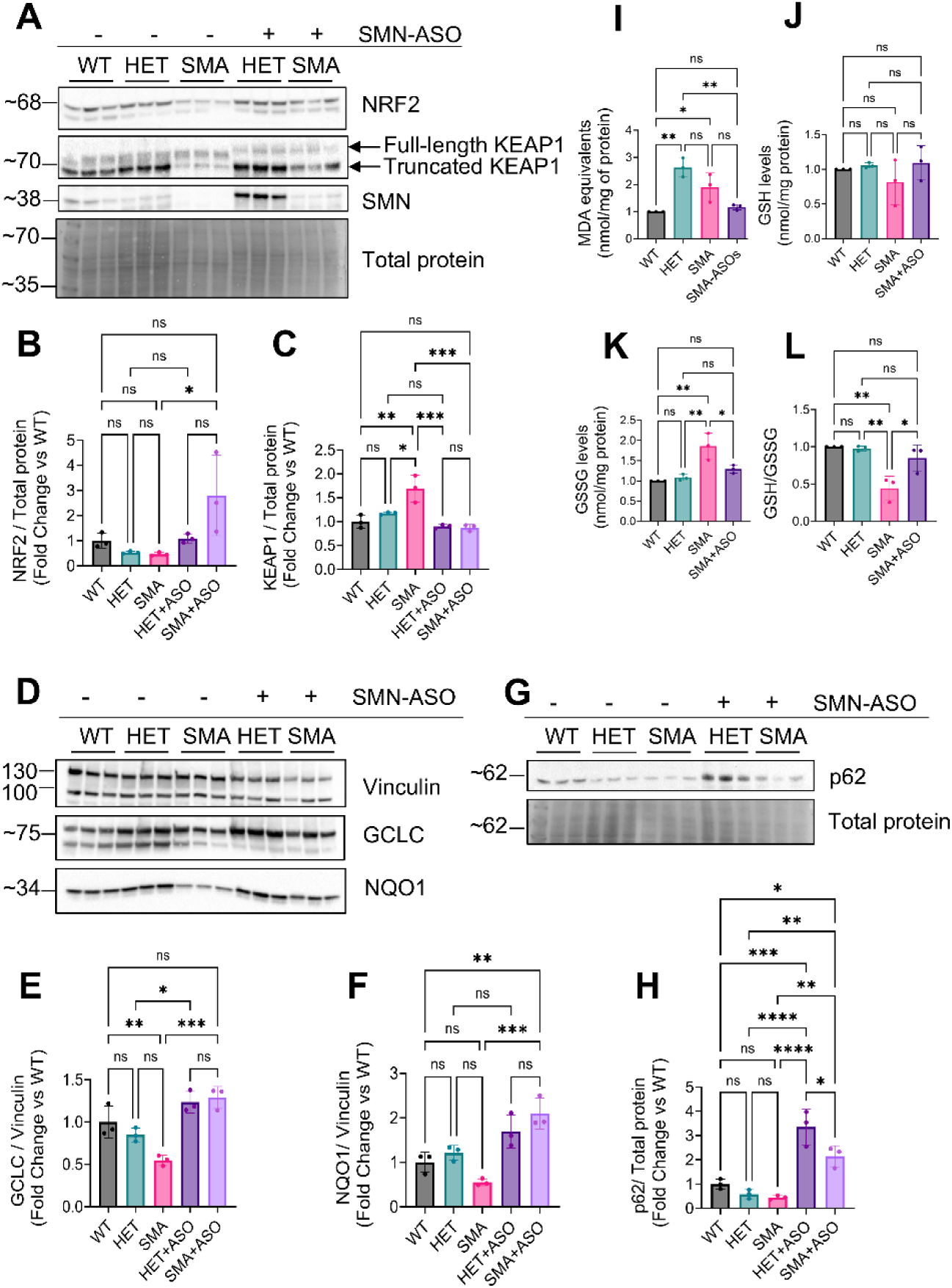
SMN-ASO restores NRF2 antioxidant signaling and hepatic redox homeostasis in SMA liver. (A) Representative Western blots of NRF2, KEAP1 (full-length and truncated forms), and SMN in liver lysates from WT, HET, SMA, HET+ASO, and SMA+ASO mice at postnatal day 10 (P10). Total protein staining was used as a loading control. (B–C) Densitometric quantification of NRF2 (B) and full-length KEAP1 (C) protein levels normalized to total protein and expressed as fold change relative to WT. Untreated SMA liver shows reduced NRF2 and increased KEAP1 levels. SMN-ASO treatment increases NRF2 abundance and reduces KEAP1 toward control levels in SMA, whereas ASO administration does not significantly alter NRF2 or KEAP1 levels in HET animals. (D) Representative Western blots of the NRF2 downstream antioxidant targets GCLC and NQO1 in liver lysates from all experimental groups. Vinculin was used as a loading control. (E-F) Quantification of GCLC (E) and NQO1 (F) protein levels normalized to Vinculin and expressed as fold change relative to WT. Both proteins are significantly reduced in SMA liver and restored to control levels following SMN-ASO treatment. (G) Representative Western blot of the autophagy adaptor protein p62 in liver lysates from WT, HET, SMA, HET+ASO, and SMA+ASO mice. Total protein staining was used as a loading control. (H) Densitometric quantification of SQSTM1/p62 protein levels normalized to total protein and expressed as fold change relative to WT. SQSTM1/p62 levels are reduced in untreated SMA liver and increased in both HET+ASO and SMA+ASO animals. (I) Malondialdehyde (MDA) levels in liver tissue, shown as a measure of lipid peroxidation. Data are expressed as MDA equivalents (nmol per mg protein). (J) Total glutathione (GSH) levels in liver tissue, expressed as nmol per mg protein. (K) Oxidized glutathione (GSSG) levels in liver tissue, expressed as nmol per mg protein. (L) GSH/GSSG ratio, which is significantly reduced in SMA and restored to control values following SMN-ASO administration. For lipid peroxidation measurements (I), data represent at least three biological replicates per group obtained from at least two independent experiments. For glutathione measurements (J-L), data represent at least three biological replicates per group obtained from at least three independent experiments. Each dot represents an independent biological replicate. Data are presented as mean ± SD. Statistical analysis was performed using one-way ANOVA followed by Tukey’s multiple-comparisons test. Significance is indicated as p < 0.05 (*), p < 0.01 (**), p < 0.001 (***), and p < 0.0001 (****); ns denotes not significant.

Analysis of canonical NRF2 transcriptional targets further supported reactivation of antioxidant signaling following SMN-ASO treatment. The glutathione synthesis enzyme GCLC and the detoxification enzyme NQO1, both significantly reduced in untreated SMA liver, were restored to levels comparable to controls in SMA+ASO animals (Fig. 9D-F), consistent with enhanced NRF2 pathway activity.

The autophagy adaptor protein SQSTM1/p62 represents an additional regulatory component that may contribute to modulation of the NRF2 pathway. SQSTM1/p62 contains a KEAP1interacting region that has been shown to sequester KEAP1 and limit KEAP1-mediated degradation when SQSTM1/p62 levels are elevated [68, 69]. In our dataset, SQSTM1/p62 abundance was reduced in untreated SMA liver but increased in both HET+ASO and SMA+ASO animals (Fig. 9G, H). This increase coincided with reduced KEAP1 levels and restoration of NRF2 and its downstream targets, suggesting that SMN-ASO may contribute to NRF2 signaling, at least in part, through mechanisms involving SQSTM1/p62-dependent regulation of KEAP1. This interpretation is consistent with the noncanonical, cysteine-independent SQSTM1/p62-KEAP1-NRF2 pathway described previously [68, 70], in which elevated SQSTM1/p62 promotes KEAP1 inactivation and antioxidant gene expression. To assess whether restoration of NRF2 signaling was accompanied by improvements in hepatic redox status, biochemical markers of lipid peroxidation and glutathione redox balance were determined (Fig. 9I-L). MDA levels were elevated in SMA liver relative to WT controls, while HET animals displayed an even greater increase compared to WT, suggesting that partial SMN insufficiency is sufficient to enhance lipid peroxidation and highlighting that HET liver is not metabolically neutral with respect to oxidative stress (Fig. 9I). In contrast, SMA+ASO mice showed MDA levels comparable to WT and HET controls, consistent with reduced lipid peroxidation following SMN restoration (Fig. 9I).

Total glutathione (GSH) levels did not differ significantly among groups (Fig. 9J); however, oxidized glutathione (GSSG) was markedly increased in untreated SMA liver and was normalized by SMN-ASO treatment (Fig. 9K). Consequently, the GSH/GSSG ratio was significantly reduced in SMA and restored to control values in SMA+ASO mice (Fig. 9L). These changes parallel the restoration of NRF2 pathway activity and are consistent with improved redox buffering capacity following SMN-ASO treatment. Notably, SMN-ASO treatment increased GPX4 protein abundance in both HET+ASO and SMA+ASO livers to levels exceeding those observed in untreated WT controls (Fig.S12), consistent with a heightened antioxidant response associated with SMN restoration

Collectively, these data indicate that partial SMN restoration is associated with normalization of NRF2-dependent antioxidant signaling and improved hepatic redox balance in symptomatic SMA liver.

## Discussion

In this study we performed the first unbiased proteomic profiling of liver tissue in the Taiwanese SMA mouse model at the symptomatic stage P10. This dataset allowed us to define the major molecular pathways affected by SMN deficiency in the liver and positioned them within a broader metabolic and mitochondrial context. As summarized in Figure 10, symptomatic SMA liver exhibits robust activation of the heme-biosynthetic pathway together with selective mitochondrial iron accumulation, both of which can promote excessive ROS. In parallel, we identified selective destabilization of the iron–sulfur cluster-containing Complex II, and early down-regulation of PGC1α at P5. Whereas the NRF2 axis initially mounted a compensatory antioxidant response at P5, as evidenced by upregulating NQO1 and GCLC, this response collapsed by P10, resulting in impaired NRF2-regulated antioxidant defenses. Early SMN-ASO administration at P1 prevented most abnormalities observed at P10. Mechanistically, increased KEAP1 and reduced SQSTM1/p62 levels in SMA liver explain the failure of NRF2 activation; ASO treatment restored NRF2 via a SQSTM1/p62-KEAP1-dependent, noncanonical NRF2 activation mechanism.

**Figure 10.**
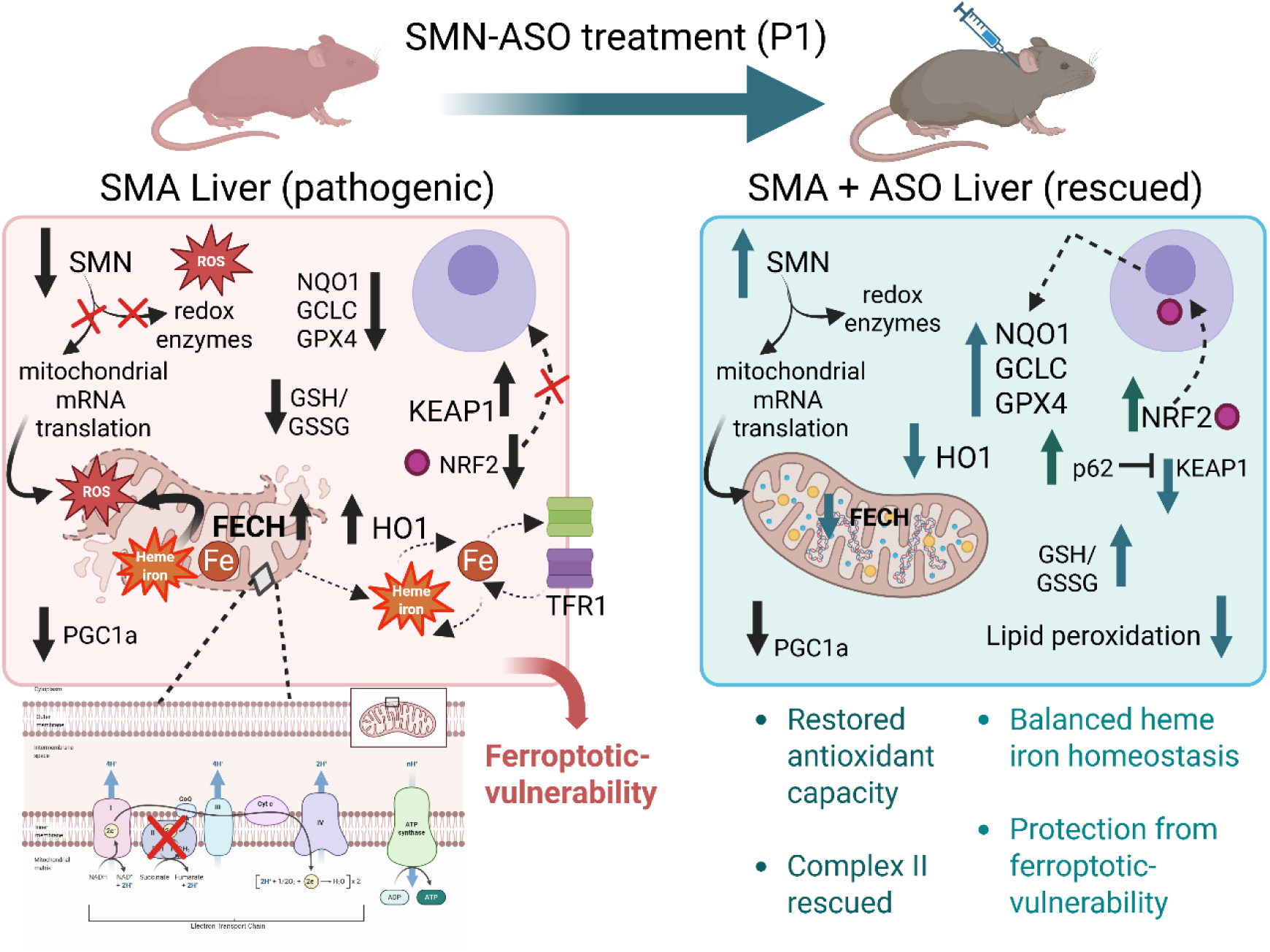
Integrated model of heme-iron-mitochondrial dysfunction and NRF2 axis dysregulation in SMA liver and its modulation by SMN-ASO. Schematic summary of molecular and metabolic alterations identified in SMA liver at the symptomatic stage (P10) and their modulation by early SMN-ASO treatment. In untreated SMA liver (left), SMN deficiency is associated with activation of the heme-biosynthetic pathway, increased ferrochelatase (FECH) abundance, increased heme catabolism pathways (e.g. HO1), and selective accumulation of mitochondrial hemic and non-hemic iron. This iron burden, together with altered iron-sulfur (Fe-S) cluster homeostasis, promotes reactive oxygen species (ROS) generation and selective destabilization of the Fe-S-containing respiratory Complex II. In parallel, mitochondrial biogenesis is impaired, as indicated by reduced PGC1α levels detectable already at the early symptomatic stage (P5). At P10, SMA liver exhibits repression of NRF2-dependent antioxidant signaling, characterized by increased KEAP1 abundance, reduced SQSTM1/p62 levels, and decreased expression of NRF2 target genes involved in glutathione synthesis and redox detoxification (e.g., NQO1, GCLC, GPX4). The resulting redox imbalance, together with mitochondrial iron loading, favors lipid peroxidation and ferroptosis susceptibility. Early postnatal SMN-ASO treatment (right) prevents mitochondrial iron accumulation, restores Complex II integrity and activity, and normalizes redox homeostasis. SMN-ASO reduces KEAP1 abundance, increases SQSTM1/p62 levels, stabilizes NRF2, and reactivates NRF2-dependent antioxidant programs through a noncanonical SQSTM1/p62-KEAP1-NRF2 mechanism.

A striking and unexpected finding was the marked upregulation of FECH, at P5 and P10. Although hemoprotein alterations and HO1 induction have been previously reported in *Smn2B/−* mice [67], the enzymatic machinery of heme synthesis itself had not been implicated. FECH elevation suggests increased mitochondrial heme synthetic capacity and flux through the terminal step of Fe_2+_ insertion [55], consistent with coordinated increases in hepatic HO1, hemopexin, haptoglobin, and BLVRB [57, 71–73]. These data support a model in which SMA liver experiences elevated heme burden and compensatory heme turnover activation. Importantly, we refine past reports of hepatic iron accumulation [17, 66, 67], by demonstrating that excess iron is predominantly mitochondrial and present in both hemic and non-hemic forms, directly linking heme metabolism to mitochondrial iron loading.

Downregulation of PGC1α, as early as at P5, when FECH is already markedly increased, combined with absent proteomic ALAS1 detection, argues against the canonical PGC1αALAS1 axis [74] as the main driver of the heme changes observed in SMA liver. Instead, our data suggest mitochondrial compensation for rising intramitochondrial iron levels by upregulating FECH to channel excess iron into heme [57]. However, reduced PGC1α is expected to limit mitochondrial biogenesis and assembly of heme-requiring respiratory complexes [75, 76]. This mismatch between heme production and heme utilization likely triggers the induction of heme-turnover pathways and provides a mechanistic basis for the selective vulnerability of Complex II, whose assembly requires coordinated heme and Fe-S cluster maturation [61, 77].

The identification of selective Complex II impairment offers a mechanistic bridge between these disrupted heme/iron metabolism and mitochondrial dysfunction. Complex II links the Krebs cycle to the electron transport chain and requires multiple [Fe-S] clusters and a heme b group for stability [61, 78]. Reduced SDHB levels and loss of Complex II activity reflect destabilization of this Fe–S/heme-dependent complex. Unlike Complexes I or IV, which remained structurally and functionally intact, Complex II appears intrinsically vulnerable to combined disturbances in Fe-S cluster maturation, heme incorporation, and iron homeostasis. Correspondingly, proteomics revealed increased ISCA2 and ABCB-related components of FeS maturation, but decreased ferredoxin proteins (FDXR, FDX1), essential for both Fe-S cluster and heme maturation, an imbalance consistent with redox strain and compensatory yet incomplete mitochondrial biogenesis responses.

At P10, collapse of NRF2 signaling further exacerbates this metabolic and oxidative stress. Reduced NRF2, elevated KEAP1, and downregulation of NQO1, GCLC, and GPX4 indicate impaired antioxidant defense. NRF2’s role in iron/ heme metabolism [79], and redox buffering explained why its suppression exacerbates mitochondrial iron-driven oxidative stress. Early GPX4 reduction at P5 suggests that ferroptosis-related vulnerability precedes the NRF2 collapse, aligning with increased TBARS and the broader literature linking SMN deficiency to GPX family alterations [12].

SMN-ASO treatment clarified which defects arise directly from SMN deficiency versus secondary consequences. FECH overexpression, mitochondrial iron overload, Complex II impairment, NRF2 suppression, and KEAP1 elevation were all selectively normalized in SMA treated mice. Complete correction of mitochondrial hemic and non-hemic iron indicates that iron overload is a reversible SMN-dependent phenotype, not secondary tissue damage. Full rescue of Complex II substantiates a functional connection between Fe-S/heme disturbances and respiratory impairment. However, although SMN-ASO treatment significantly reduced FECH abundance in SMA liver, FECH levels remained elevated relative to WT controls, indicating partial rather than complete normalization of heme biosynthetic capacity.

In contrast, GPX4 and SQSTM1/p62 reflected SMN dosage sensitivity rather than SMA-specific pathology. Both increased after ASO treatment in SMA and HET mice, revealing graded SMN-dependent regulation. Given SQSTM1/p62’s roles in autophagy and KEAP1 repression [80], these findings provide a mechanistic basis for ASO-mediated NRF2 restoration and extend the known SMA-associated autophagy disturbances [81–84] to the liver.

Together, our findings support a unifying model: SMN deficiency perturbs heme synthesis, FeS cluster maturation, and mitochondrial iron allocation, selectively impairing Complex II and triggering a cascade of oxidative and ferroptotic vulnerability driven by NRF2 suppression and GPX4 loss. The SMA liver emerges as a metabolically fragile organ undergoing coordinated mitochondrial-heme-iron-redox remodeling that is SMN-dependent and largely reversible. These hepatic defects likely contribute to systemic metabolic fragility in SMA, although tissuespecific metabolic context may shape their manifestation. Because SMN-ASO administration was systemic, our data cannot distinguish liver-autonomous from whole-body effects.

Finally, these findings have translational implications. Partial persistence of altered PGC1α and FECH after SMN restoration, suggests that early-life mitochondria and heme metabolic defects may endure despite normalized redox status, highlighting the importance of therapeutic timing. The identification of mitochondrial iron overload, Complex II impairment, NRF2 suppression, and GPX4 deficiency suggests rational combinatorial therapies including ferroptosis inhibition, NRF2 activation, iron modulation, or mitochondrial-targeted antioxidants. The reversibility of most abnormalities with modest SMN increase indicates that SMA-associated metabolic and redox dysregulation is therapeutically pliable, not a fixed degenerative endpoint.

## Data Availability

The mass spectrometry proteomics data generated in this study have been deposited to the ProteomeXchange Consortium via the PRIDE partner repository under the dataset identifier PXD070887. Protein-level quantitative proteomics data, differential expression analyses, and enrichment results are provided as Supplementary Tables accompanying this preprint.

## CRediT authorship contribution statement

**Sofia Vrettou:** Conceptualization, Methodology, Software, Visualization, Validation, Writing, Original draft preparation. **Stefan Müller:** Methodology, Software, Data Curation, Resources, Writing, Funding Acquisition. **Brunhilde Wirth:** Conceptualization, Supervision, Writing-Reviewing and Editing, Funding Acquisition.

## Acknowledgments

We thank IONIS Pharmaceuticals for providing the SMN-ASOs. We thank Roman Rombo for technical assistance in maintaining, breeding and treating the mice. This work was supported by the large instrument grant INST 216/1163-1 FUGG by the German Research Foundation (DFG Großgeräteantrag) to SM, European Union’s Horizon 2020 Marie Skłodowska-Curie Program (project 956185; SMABEYOND) to BW and the Center for Molecular Medicine Cologne (project C18) to BW. The figures 1A, 3A, 4E, 5C, 7A, 8A, 10 were created using BioRender.com.

## Abbreviations

SMA: Spinal muscular atrophy
SMN: Survival motor neuron
ASO: Antisense oligonucleotide
WT: Wild type
HET: Heterozygous
P5 / P10: Postnatal day 5 / postnatal day 10
OXPHOS: Oxidative phosphorylation
BN-PAGE: Blue native polyacrylamide gel electrophoresis
CN-PAGE: Clear native polyacrylamide gel electrophoresis
FECH: Ferrochelatase
HMBS: Hydroxymethylbilane synthase
ALAD: Aminolevulinate dehydratase
HO-1: Heme oxygenase 1
NRF2: Nuclear factor erythroid 2–related factor 2
KEAP1: Kelch-like ECH-associated protein 1
GPX4: Glutathione peroxidase 4
NQO1: NAD(P)H quinone dehydrogenase 1
GCLC: Glutamate-cysteine ligase catalytic subunit
GSH: Reduced glutathione
GSSG: Oxidized glutathione
MDA: Malondialdehyde
Fe-S: Iron-sulfur
PGC1α: Peroxisome proliferator-activated receptor gamma coactivator 1-alpha

## Supplementary material

Supplementary Figures S1-S12

Supplementary Figure 13: Uncropped Western blots and native gels

Supplementary Table S1: Antibodies & Primers

Supplementary Table S2: Complete protein identification and quantification

Supplementary Table S3: Volcano matrix WT vs SMA

Supplementary Table S4: Volcano matrix HET vs SMA

Supplementary Table S5: Differentially expressed proteins (Venn)

Supplementary Table S6: Enriched pathways WT vs SMA

Supplementary Table S7: Enriched pathways WT vs SMA

Supplementary Table S8: ClueGO results

