## Supplementary Figures S1-S12 for "SMN deficiency disrupts hepatic mitochondrial iron homeostasis and NRF2-dependent redox control in spinal muscular atrophy"

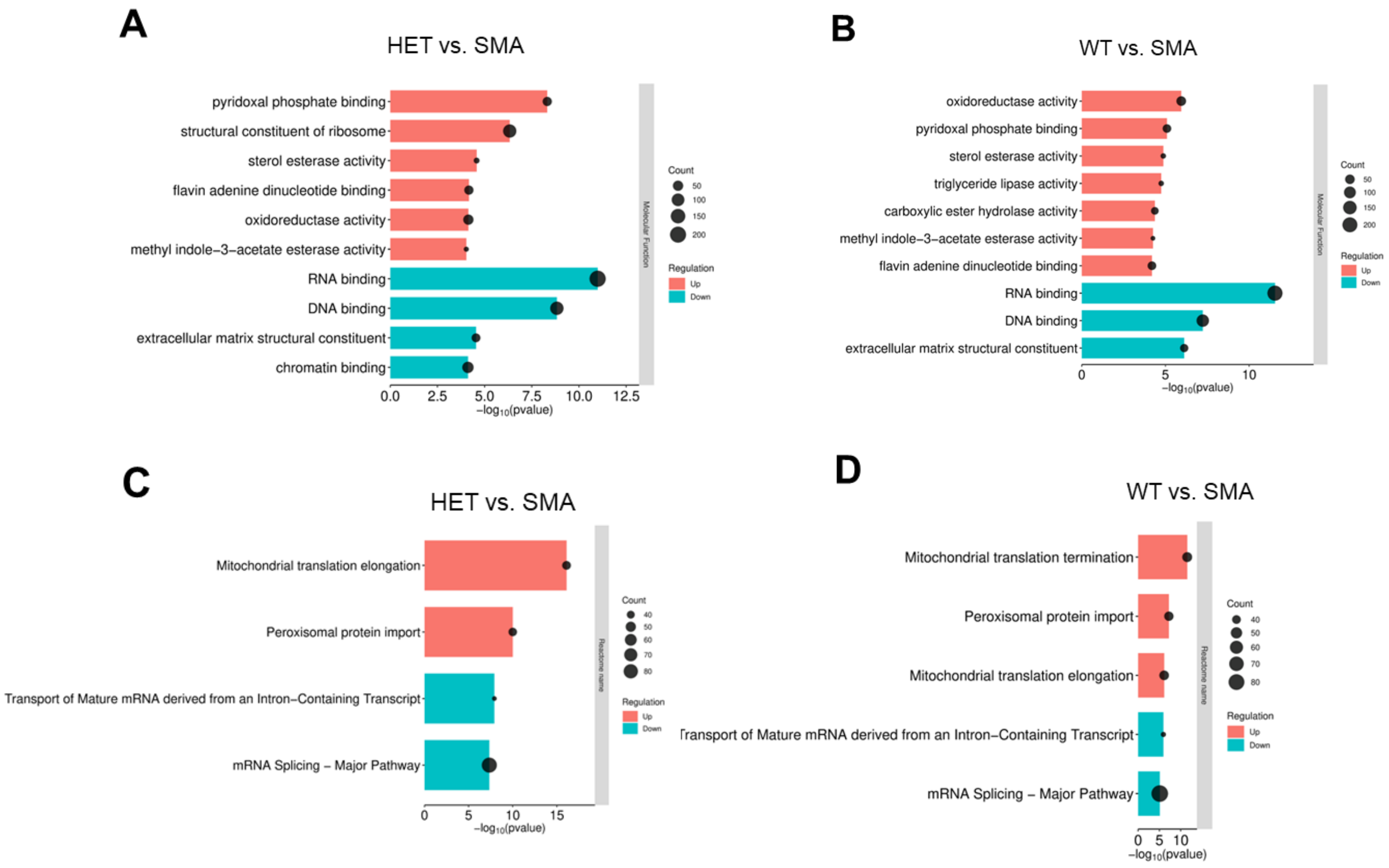


**Supplementary Figure S1: Molecular function and reactome pathway enrichment in SMA liver.**

**(A–B)** Gene ontology molecular function (MF) enrichment analysis comparing HET vs. SMA (A) and WT vs. SMA (B) liver proteomes. Bars represent significantly enriched MF terms derived from 1D enrichment analysis in Perseus, visualized as –log10(p-values). Terms enriched among proteins increased in SMA (red) include oxidoreductase activity, pyridoxal phosphate binding, and sterol/ester hydrolase activity, whereas terms enriched among proteins decreased in SMA (blue) are predominantly associated with RNA binding, DNA binding, chromatin binding, and extracellular matrix structural constituents. Dot size indicates the number of proteins associated with each term.
**(C–D)** Reactome pathway enrichment analysis for HET vs. SMA (C) and WT vs. SMA (D). Pathways enriched among proteins increased in SMA (red) are centered on mitochondrial translation elongation/termination and peroxisomal protein import, while pathways enriched among proteins decreased in SMA (blue) include mRNA splicing and transport of mature mRNA derived from intron-containing transcripts. Enrichment significance is shown as –log10(p-values), with dot size reflecting pathway size.

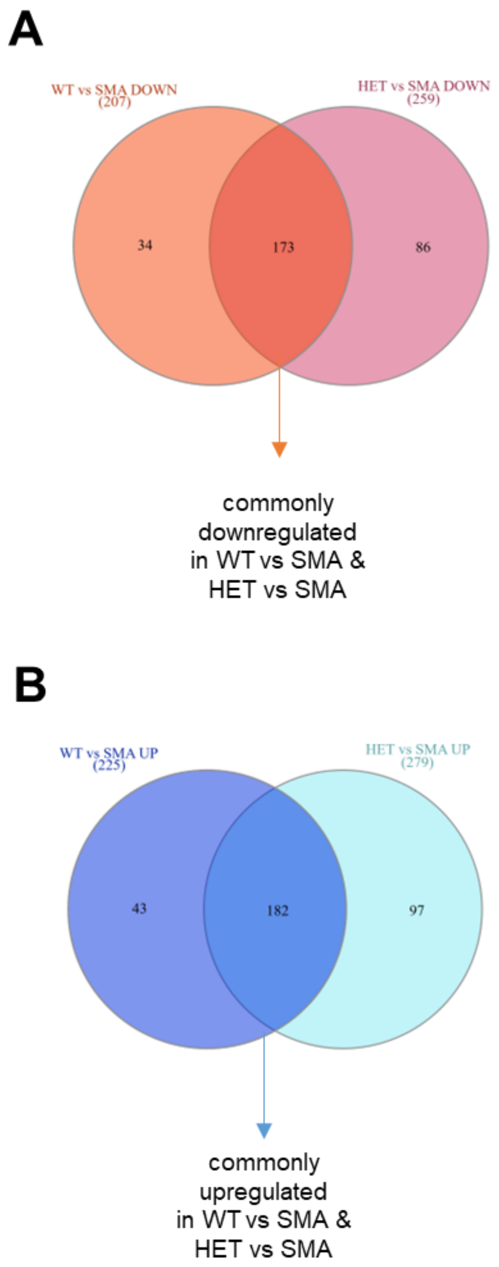


**Supplementary Figure S2: Overlap of differentially abundant proteins between WT vs. SMA and HET vs. SMA comparisons.**

Venn diagrams illustrating the overlap and comparison-specificity of proteins downregulated (A) and upregulated (B) in SMA liver relative to WT and HET controls.
**(A)** A total of 173 proteins were commonly downregulated in SMA in both WT vs. SMA and HET vs. SMA comparisons, with 34 proteins uniquely decreased in WT vs. SMA and 86 uniquely decreased in HET vs. SMA.
**(B)** A total of 182 proteins were commonly upregulated in SMA in both comparisons, with 43 proteins uniquely increased in WT vs. SMA and 97 uniquely increased in HET vs. SMA. The complete lists of proteins corresponding to each Venn diagram sector, including identifiers and statistical parameters, are provided in the Supplementary Table S2.

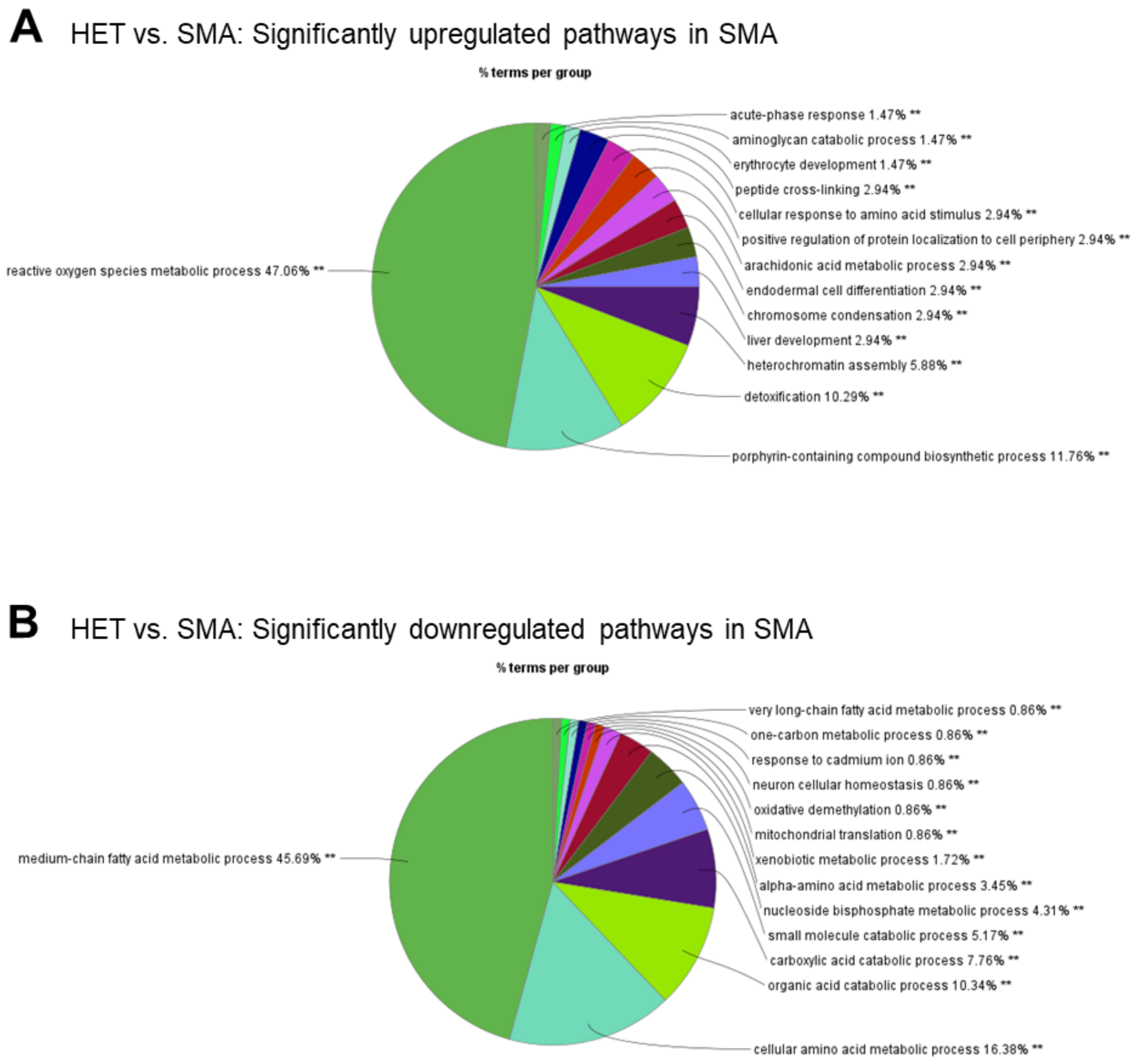


**Supplementary Figure S3. Functional enrichment of proteins differentially expressed between SMA and HET liver.**

Functional enrichment and clustering of proteins differentially expressed in SMA relative to HET were performed using STRING-derived interaction networks visualized in Cytoscape and grouped by ClueGO based on shared biological processes.
**(A)** Pie chart showing the relative distribution of significantly enriched biological process clusters among proteins upregulated in SMA, highlighting dominant themes related to reactive oxygen species metabolism, detoxification pathways, and porphyrin-containing compound biosynthesis, alongside additional metabolic and chromatin-associated processes.
**(B)** Pie chart depicting the relative contribution of enriched biological process clusters among proteins downregulated in SMA, revealing a predominance of pathways linked to amino acid and fatty acid metabolism, organic and carboxylic acid catabolism, and mitochondrial translational and metabolic functions.
Detailed term lists, enrichment statistics, and cluster assignments are provided in the corresponding Supplementary Table S3.

**A** WT vs. SMA: Significantly upregulated pathways in SMA

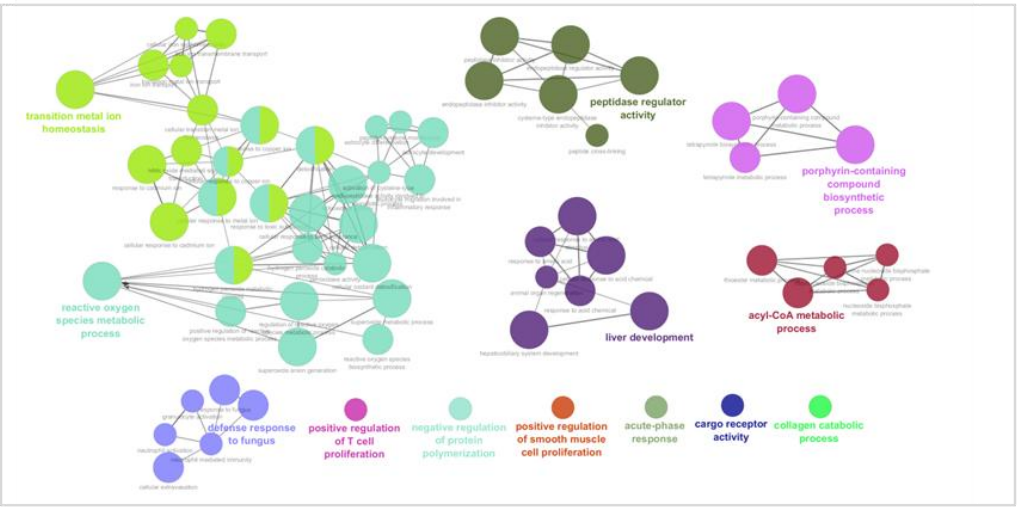


**B** WT vs. SMA: Significantly downregulated pathways in SMA

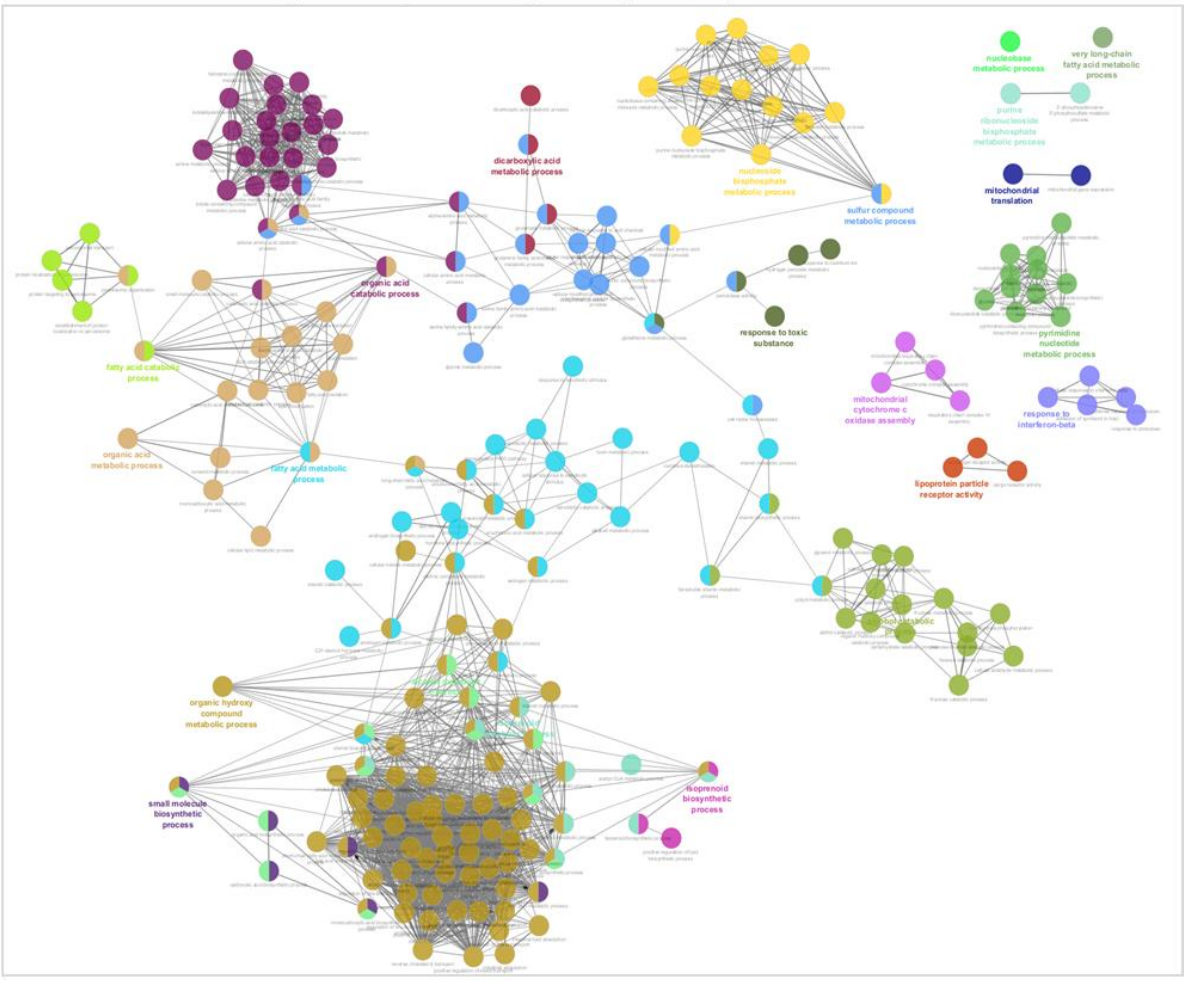


**Supplementary Figure S4. STRING-Cytoscape network analysis of proteins differentially expressed in SMA liver relative to WT control.**

**(A) Upregulated pathways in SMA (WT vs. SMA).**
Proteins significantly increased in SMA liver relative to WT were subjected to STRING interaction analysis and visualized in Cytoscape using ClueGO. Enriched biological

processes clustered predominantly around reactive oxygen species metabolism, porphyrin-containing compound and tetrapyrrole biosynthesis, metal ion homeostasis, detoxification, and stress-responsive and developmental pathways, including liver development and peptidase regulator activity. These clusters highlight a coordinated activation of redox, heme, and stress-adaptation programs in SMA liver.

**(B) Downregulated pathways in SMA (WT vs. SMA).**
Proteins significantly reduced in SMA liver relative to WT formed networks enriched for metabolic and mitochondrial processes, including fatty acid and amino acid catabolism, one-carbon metabolism, small-molecule metabolism, and mitochondrial translation. Large interconnected clusters corresponded to lipid and organic acid metabolic pathways, indicating broad suppression of core metabolic functions in SMA liver.

For both panels, nodes represent enriched biological process terms, node size reflects the percentage of mapped genes per term, and node color denotes functional group assignment according to kappa score ($\geq 0.4$). Statistical significance was defined using a Benjamini–Hochberg–corrected $FDR < 0.05$.

**A** WT vs. SMA: Significantly upregulated pathways in SMA

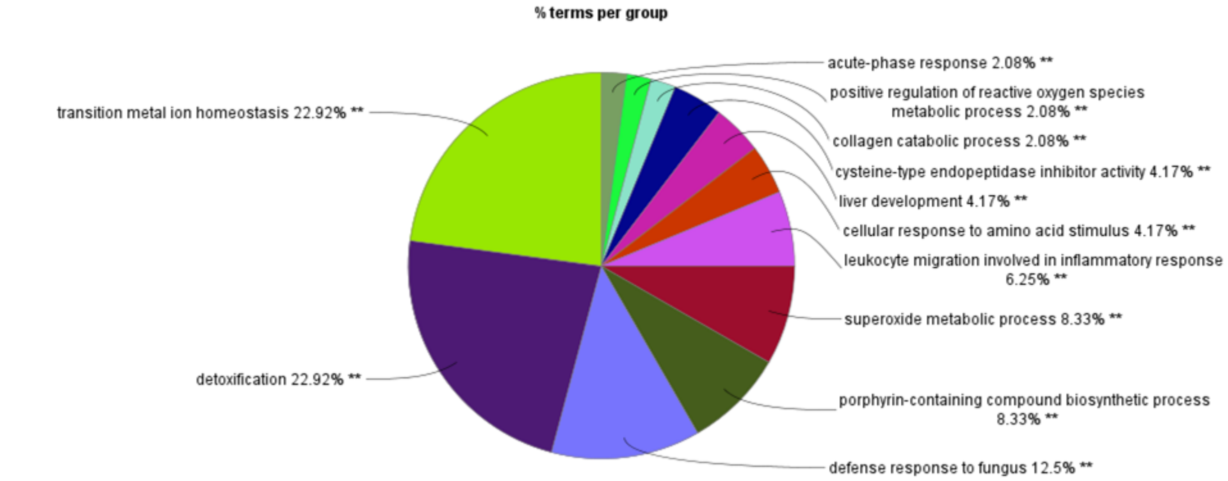


**B** WT vs. SMA: Significantly downregulated pathways in SMA

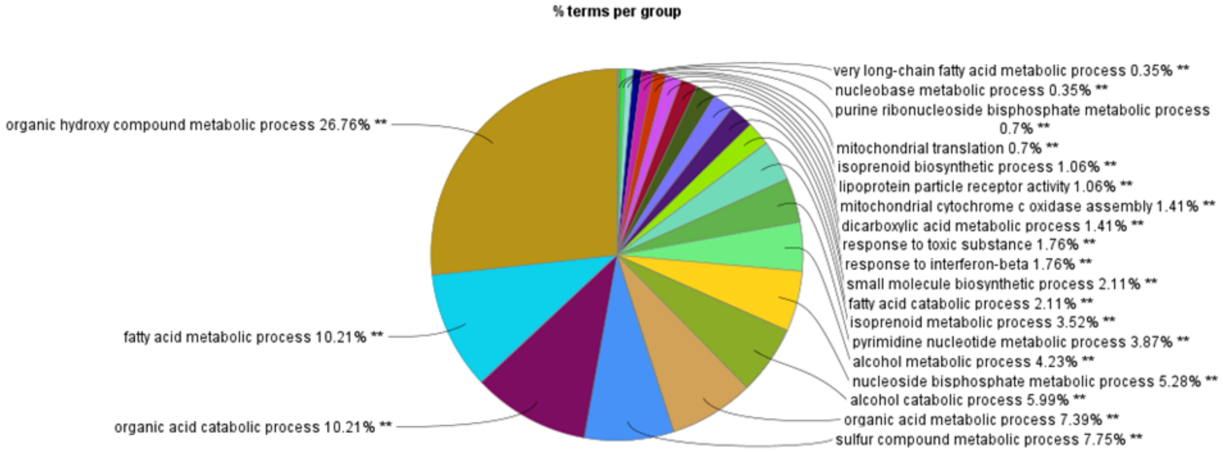


**Supplementary Figure S5. Functional enrichment of proteins differentially expressed between SMA and WT liver.**

Functional enrichment and clustering of proteins significantly altered in SMA relative to WT were performed using STRING-derived interaction networks visualized in Cytoscape and grouped by ClueGO based on shared biological processes.
**(A)** Pie chart showing the relative distribution of significantly enriched biological process clusters among proteins upregulated in SMA, highlighting dominant themes related to redox and stress responses, including reactive oxygen species metabolism, detoxification pathways, porphyrin-containing compound biosynthesis, transition metal and iron ion homeostasis, as well as immune- and inflammation-associated processes.
**(B)** Pie chart depicting the relative contribution of enriched biological process clusters among proteins downregulated in SMA, revealing a predominance of metabolic pathways, including lipid and fatty acid metabolism, small-molecule and amino-acid catabolism, cholesterol- and bile-acid-related processes, and mitochondrial translation.
Detailed term lists, enrichment statistics, and cluster assignments are provided in the corresponding Supplementary Table S3

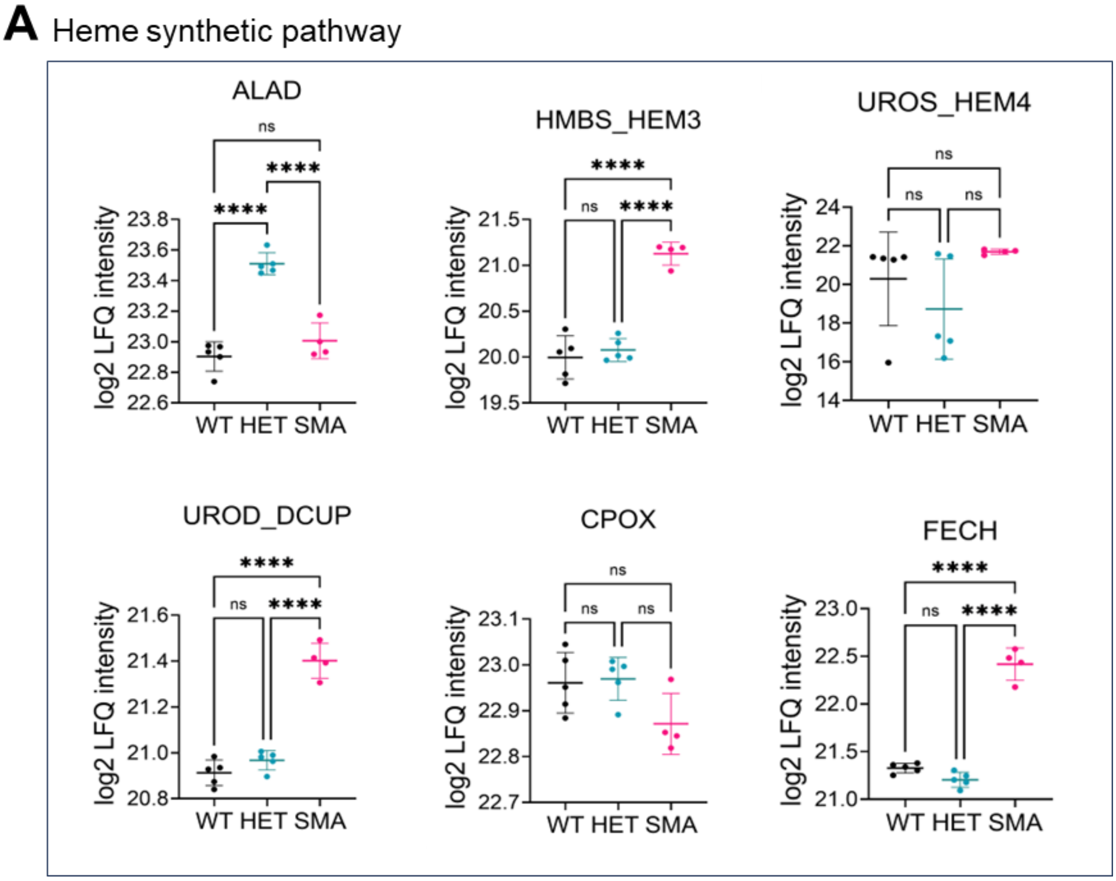


**Supplementary Figure S6. Abundance of heme biosynthesis enzymes in P10 SMA liver proteome.**

**(A)** Log2 LFQ intensities of heme biosynthesis enzymes detected in the liver proteomics dataset are shown for WT, HET, and SMA postnatal day 10 (P10) mice. The enzymes detected include Aminolevulinate dehydratase (ALAD), Hydroxymethylbilane synthase (HMBS; gene name *Hem3*), Uroporphyrinogen III synthase (UROS; gene name *Hem4*), Uroporphyrinogen decarboxylase (UROD; gene name *Urod*), Coproporphyrinogen oxidase (CPOX), and Ferrochelatase (FECH).

HMBS, UROD, and FECH showed increased abundance in SMA compared to WT and HET, reaching statistical significance as indicated. ALAD was significantly decreased in SMA compared to HET but not WT. UROS and CPOX did not show statistically significant differences across groups. Statistical analysis was performed by one-way ANOVA with multiple-comparison testing. Data are shown as mean ± SD. Asterisks denote significance levels as follows: $p < 0.05$ (*), $p < 0.01$ (**), $p < 0.001$ (***), $p < 0.0001$ (****), ns (not significant).

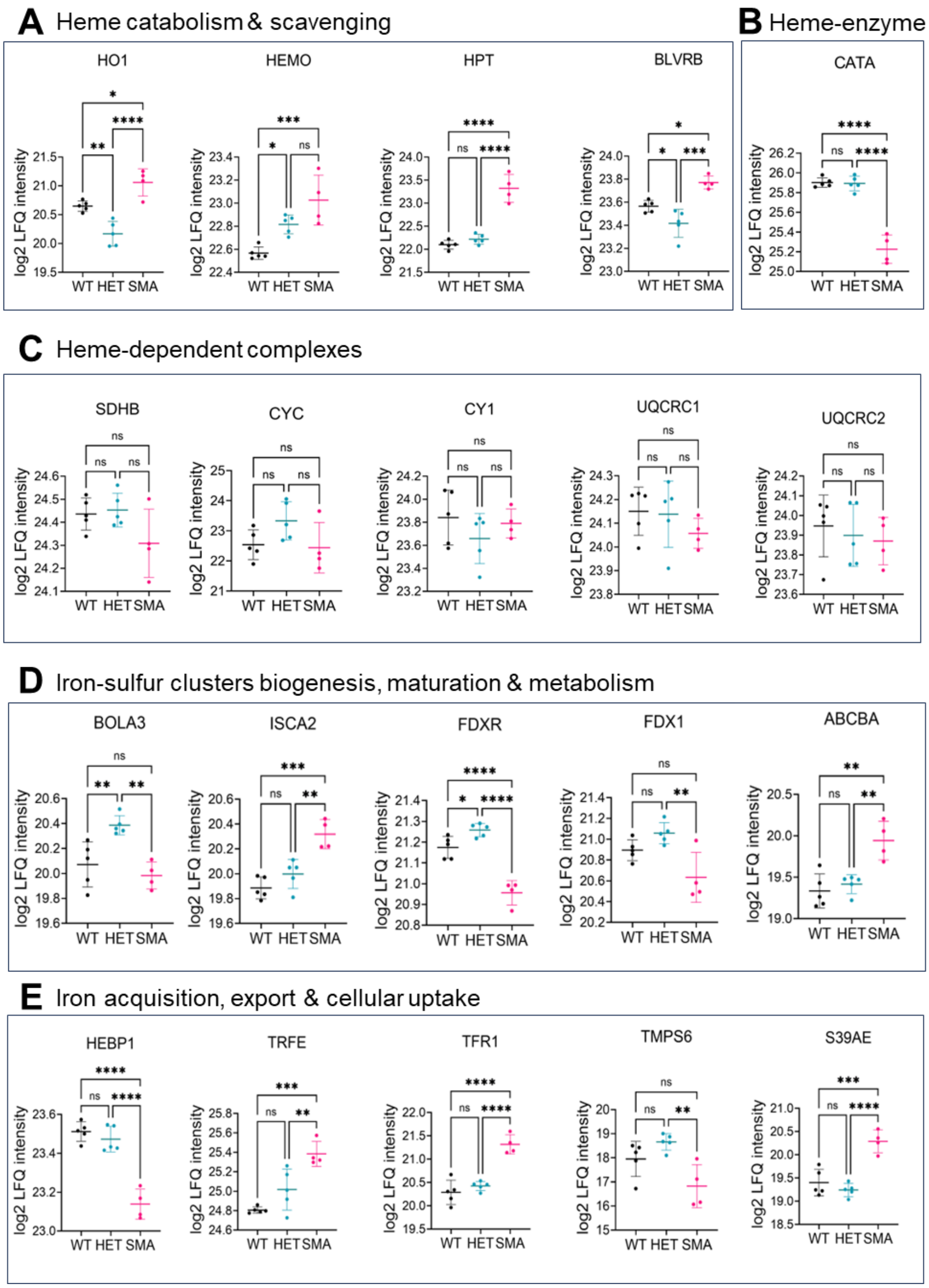


**Supplementary Figure S7. Coordinated remodeling of hepatic iron metabolism pathways in SMA revealed by proteomic profiling.**

(A) Log2 LFQ intensity profiles of proteins involved in heme catabolism and scavenging, including HO1, HEMO, HPT, and BLVRB, across WT, HET, and SMA liver samples.
(B) Log2 LFQ intensity profiles of heme-utilizing enzymes, exemplified by catalase (CAT).
(C) Log2 LFQ intensity profiles of heme-dependent mitochondrial complexes and subunits, including SDHB, CYC, CY1, UQCRC1, and UQCRC2.
(D) Log2 LFQ intensity profiles of proteins involved in iron–sulfur (Fe-S) cluster biogenesis, maturation, and metabolism, including BOLA3, ISCA2, FDXR, FDX1, and ABCBA.

(E) Log2 LFQ intensity profiles of proteins involved in iron acquisition, export, and cellular uptake, including HEBP1, TRFE, TFR1, TMPS6, and S39AE.
Each dot represents an individual biological replicate. Data were extracted from Perseus profile plots of statistically significant proteins and grouped by functional module. Together, the profiles reveal coordinated alterations in heme turnover, Fe-S cluster machinery, and iron handling in SMA liver, extending beyond heme biosynthesis to broader iron-redistribution and redox-adaptive responses.
Data are presented as individual log2 LFQ intensities. Statistical significance was determined in Perseus using Student's t-test with permutation-based FDR correction; significance is indicated as $p < 0.05$ (*), $p < 0.01$ (**), $p < 0.001$ (***), and $p < 0.0001$ (****); ns denotes not significant.

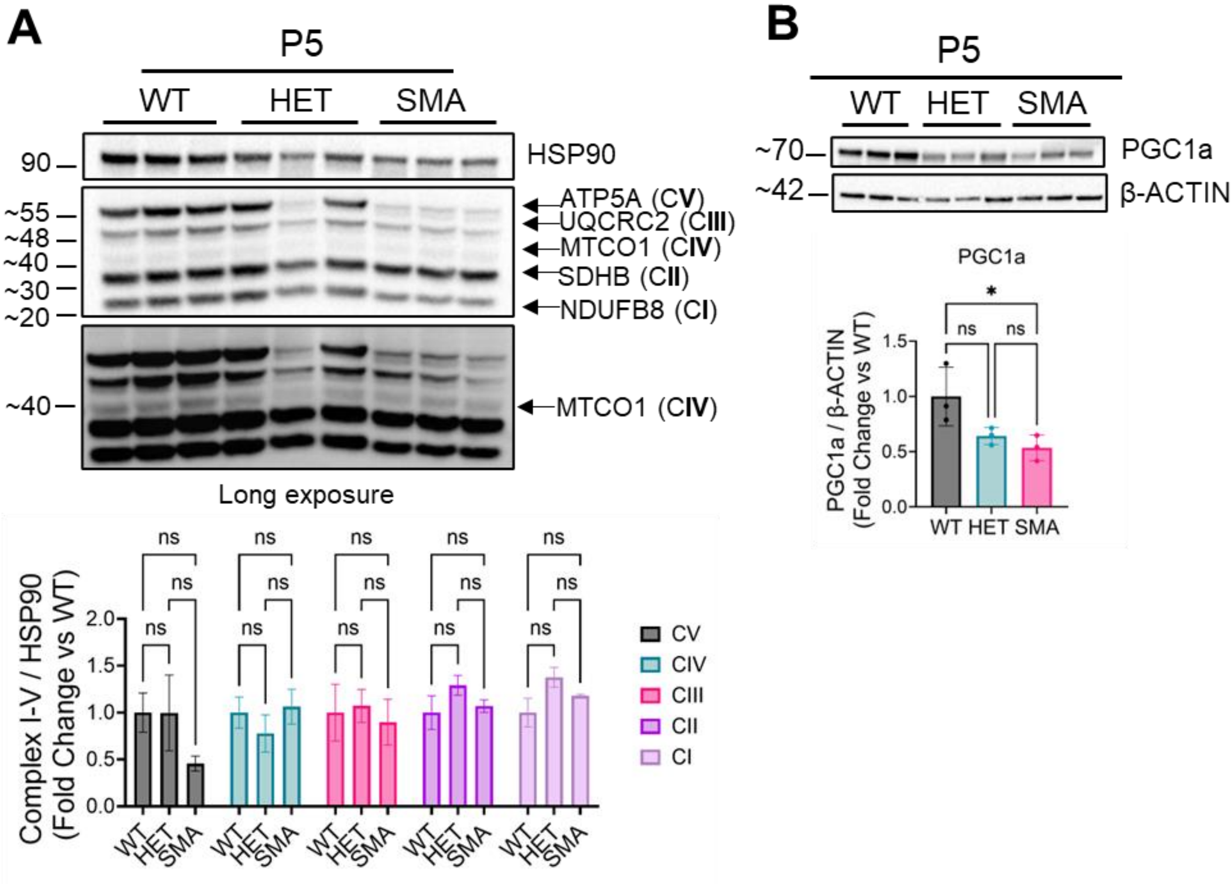


**Supplementary Figure S8. Mitochondrial OXPHOS subunits and PGC-1α levels at P5 in SMA liver.**

**(A)** Representative immunoblot of mitochondrial oxidative phosphorylation (OXPHOS) complexes I–V detected using a mitocooktail antibody in liver lysates from postnatal day 5 (P5) WT, HET, and SMA mice. Individual subunits detected correspond to Complex V (ATP5A), Complex III (UQCRC2), Complex IV (MTCO1), Complex II (SDHB), and Complex I (NDUFB8), as indicated. HSP90 was used as a loading control. A longer exposure is shown to visualize lower abundance/highly hydrophobic MTCO1 subunit.

**(B)** Representative immunoblot and densitometric quantification of PGC-1α protein levels in P5 liver lysates, normalized to β-ACTIN and expressed as fold change relative to WT.

Quantification of OXPHOS subunits revealed no significant differences across genotypes at P5, indicating preserved mitochondrial OXPHOS protein abundance at this early stage. In contrast, PGC-1α levels were significantly reduced in SMA, suggesting early impairment of mitochondrial biogenic regulation prior to the emergence of overt mitochondrial defects at later symptomatic stages. Each data represents an independent biological replicate. Data are presented as mean ± SD. Asterisks denote significance levels as follows: $p < 0.05$ (*), $p < 0.01$ (**), $p < 0.001$ (***), $p < 0.0001$ (****), ns (not significant).

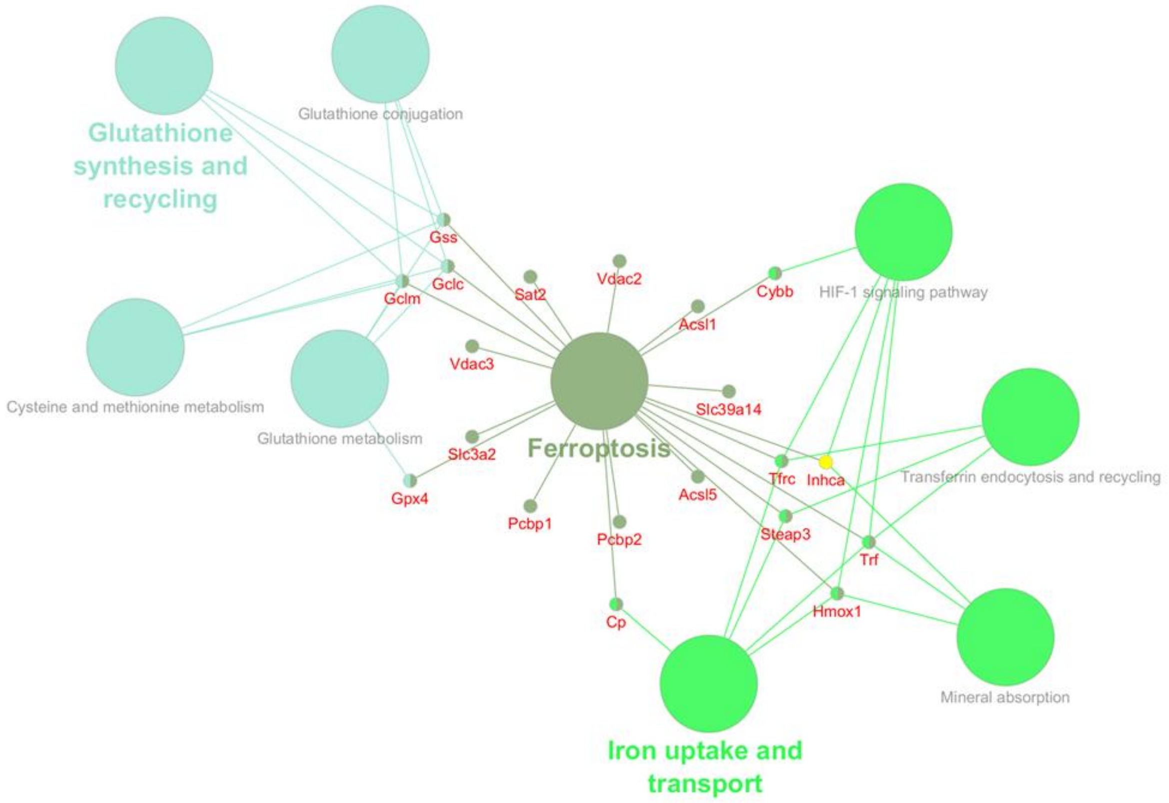


**Supplementary Figure S9. Network representation of ferroptosis-associated proteins detected in P10 SMA liver proteome.**

Proteins classified under the ferroptosis category in Perseus profile-plot analysis of the postnatal day 10 (P10) liver proteomics dataset were imported into STRING and visualized in Cytoscape, followed by functional enrichment using ClueGO with KEGG and Reactome pathway annotations. Protein nodes represent ferroptosis-associated targets detected in the dataset, while larger nodes correspond to enriched functional pathways.

The network highlights the integration of glutathione synthesis and recycling, cysteine and methionine metabolism, iron uptake and transport, transferrin endocytosis and recycling, mineral absorption, and HIF-1 signaling, illustrating the convergence of redox control, iron handling, lipid metabolism, and mitochondrial-associated processes within the ferroptosis framework.

Node labels indicate individual proteins, and edges represent known functional or pathway-based associations as curated in STRING, KEGG, and Reactome. This network summarizes the coordinated involvement of glutathione metabolism and iron homeostasis pathways among ferroptosis-associated proteins detected in P10 SMA liver proteomics.

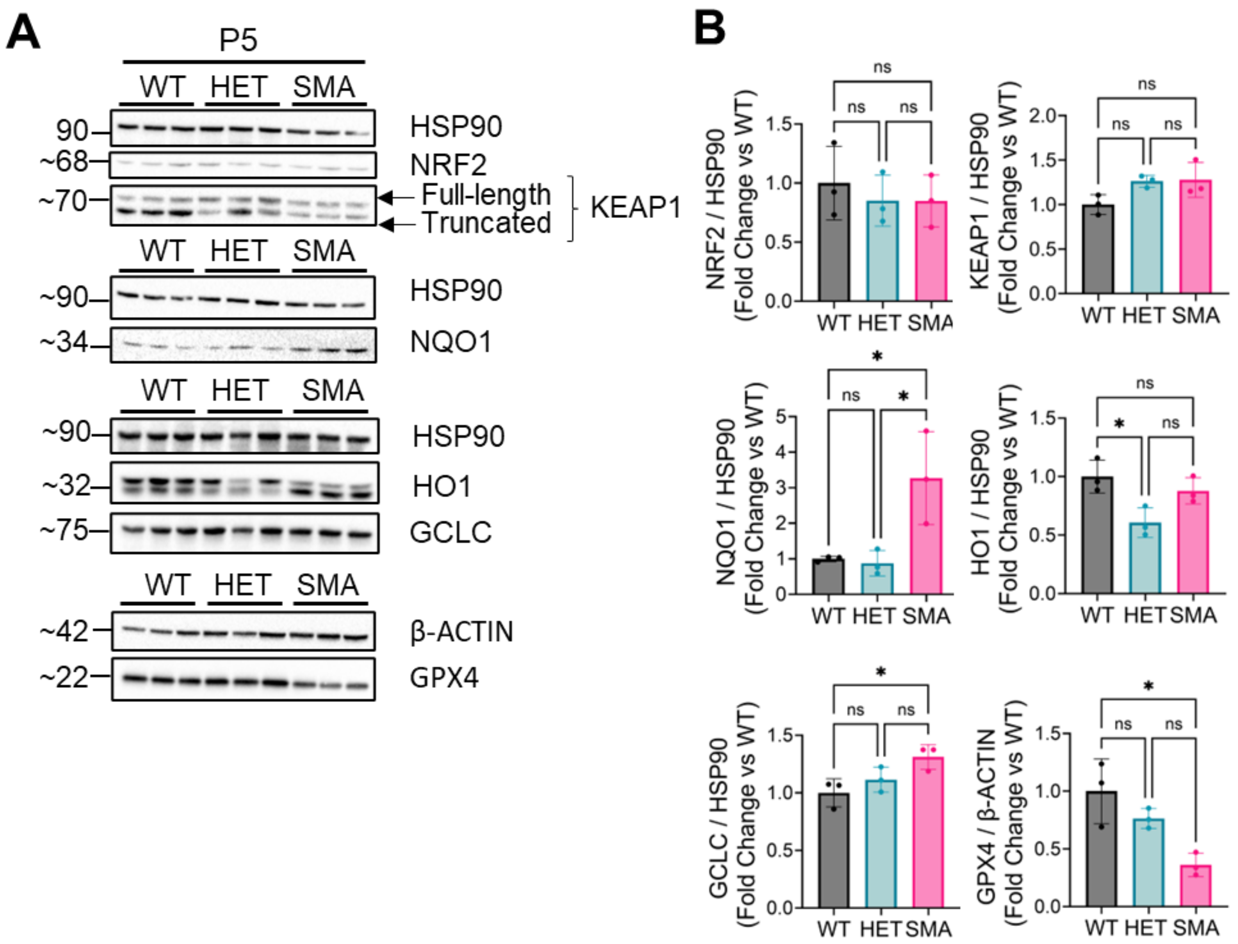


**Supplementary Figure S10. NRF2 signaling axis in P5 SMA liver.**

**(A)** Representative immunoblots showing protein levels of NRF2, KEAP1, NQO1, HO1, GCLC, and GPX4 in liver lysates from postnatal day 5 (P5) WT, HET, and SMA mice. HSP90 or β-ACTIN were used as loading controls as indicated.
**(B)** Densitometric quantification of immunoblot signals normalized to the corresponding loading control and expressed as fold change relative to WT.

At P5, NRF2 and KEAP1 protein levels were not significantly altered across genotypes. In contrast, NQO1 and GCLC were significantly increased in SMA, whereas GPX4 was significantly reduced compared to WT. HO1 levels were significantly decreased in HET compared to WT, but did not differ significantly between SMA and WT or HET. Each data represents an independent biological replicate. Data are shown as mean ± SD. Statistical analysis was performed using one-way ANOVA with appropriate multiple-comparison testing. Asterisks denote significance levels as follows: $p < 0.05$ (*), $p < 0.01$ (**), $p < 0.001$ (***), $p < 0.0001$ (****), ns (not significant).

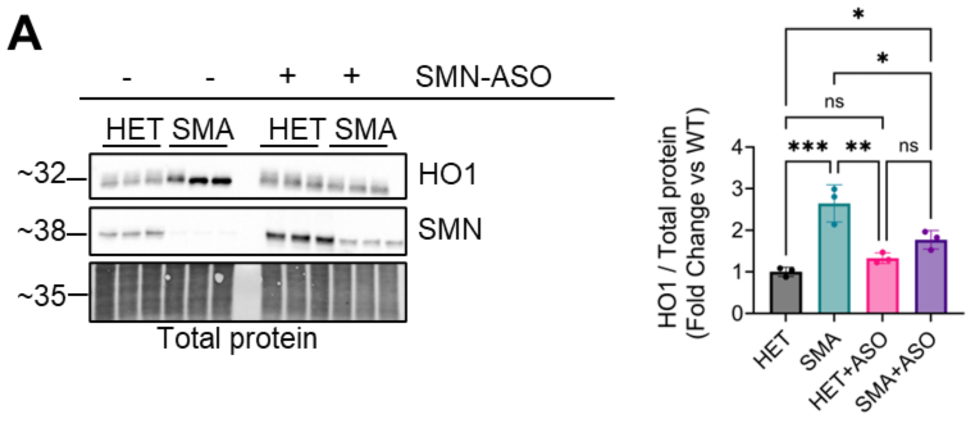


**Supplementary Figure S11. SMN-ASO treatment partially normalizes HO-1 protein levels in SMA liver.**

**(A)** Representative Western blots of whole-liver lysates from HET and SMA mice treated with vehicle (–) or SMN-targeting antisense oligonucleotides (SMN-ASO, +). HO1 and SMN protein levels are shown, with total protein staining used as the loading control.

**(B)** Quantification of HO-1 protein levels normalized to total protein and expressed as fold change relative to untreated HET controls. Each data represents an independent biological replicate. Data are shown as mean ± SD. Statistical analysis was performed using one-way ANOVA with appropriate multiple-comparison testing. Asterisks denote significance levels as follows: $p < 0.05$ (*), $p < 0.01$ (**), $p < 0.001$ (***), $p < 0.0001$ (****), ns (not significant).

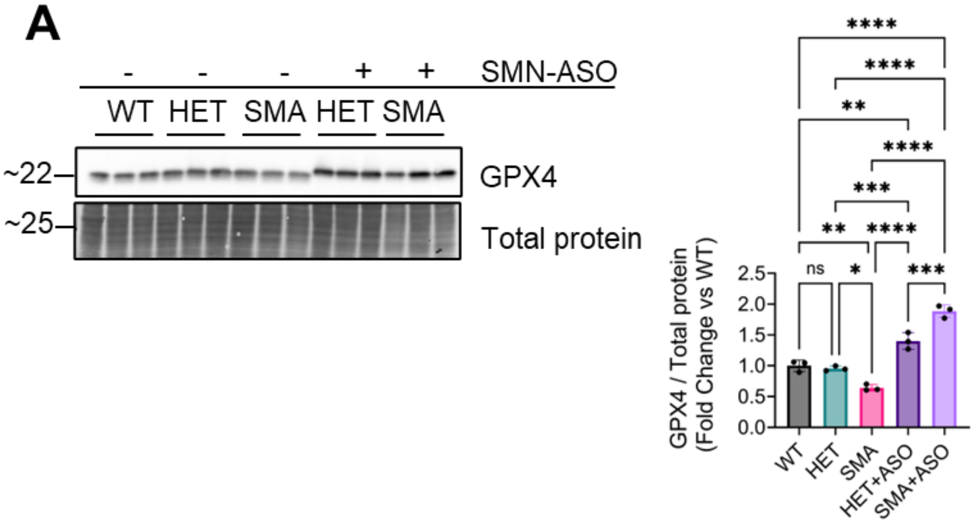


**Figure S12. SMN-ASO induces GPX4 protein expression in SMA and HET liver**

**(A)** Representative Western blot and densitometric quantification of glutathione peroxidase 4 (GPX4) protein levels in whole-liver lysates from WT, HET, SMA, HET+ASO, and SMA+ASO mice at postnatal day 10 (P10). GPX4 levels are significantly reduced in untreated SMA liver compared to WT and HET. SMN-ASO treatment markedly increases GPX4 abundance in both HET and SMA animals, reaching levels significantly higher than those observed in untreated WT and HET controls. Protein levels were normalized to total protein staining. Each data represents an independent biological replicate. Data are shown as mean ± SD. Statistical analysis was performed using one-way ANOVA with appropriate multiple-comparison testing. Asterisks denote significance levels as follows: $p < 0.05$ (*), $p < 0.01$ (**), $p < 0.001$ (***), $p < 0.0001$ (****), ns (not significant).
