## Supplementary Figures S13 for "SMN deficiency disrupts hepatic mitochondrial iron homeostasis and NRF2-dependent redox control in spinal muscular atrophy"

### Figure 4A

Samples lane order: WT-1, WT-2, WT-3, HET-1, HET-2, HET-3, SMA-1, SMA-2, SMA-3, SMA-4.

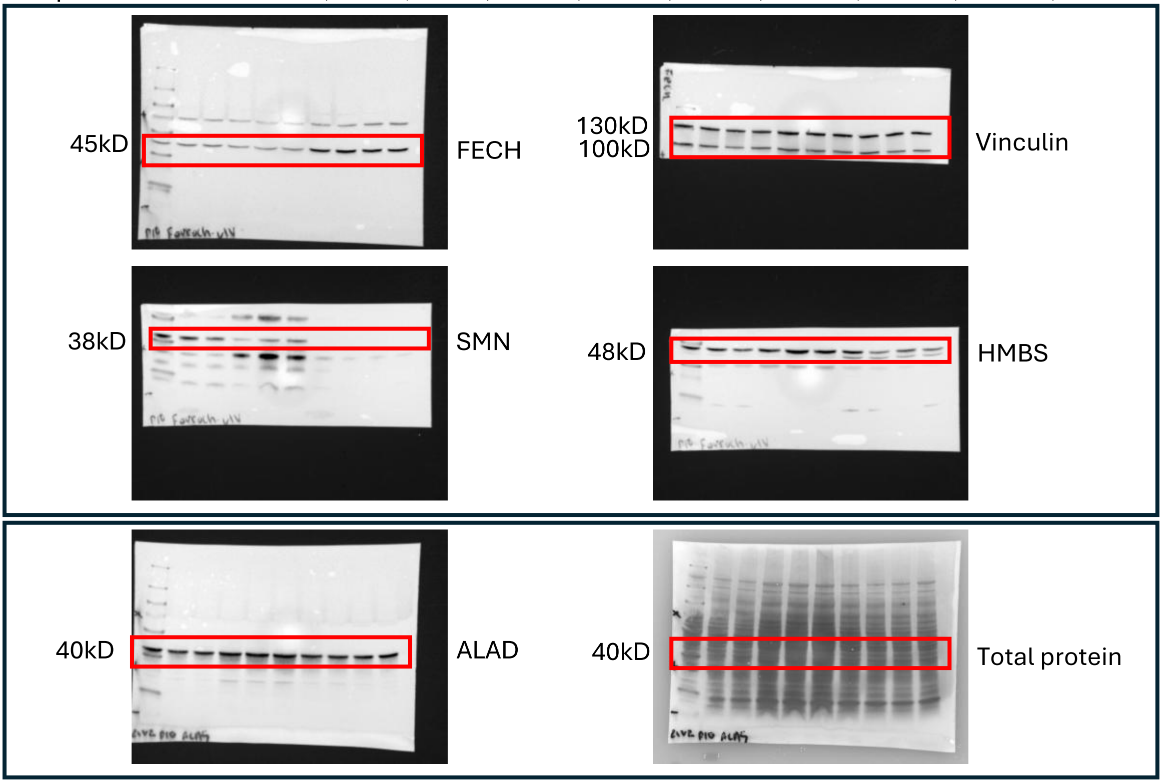


### Figure 4B

Samples lane order: WT-1, WT-2, WT-3, HET-1, HET-2, HET-3, SMA-1, SMA-2, SMA-3.

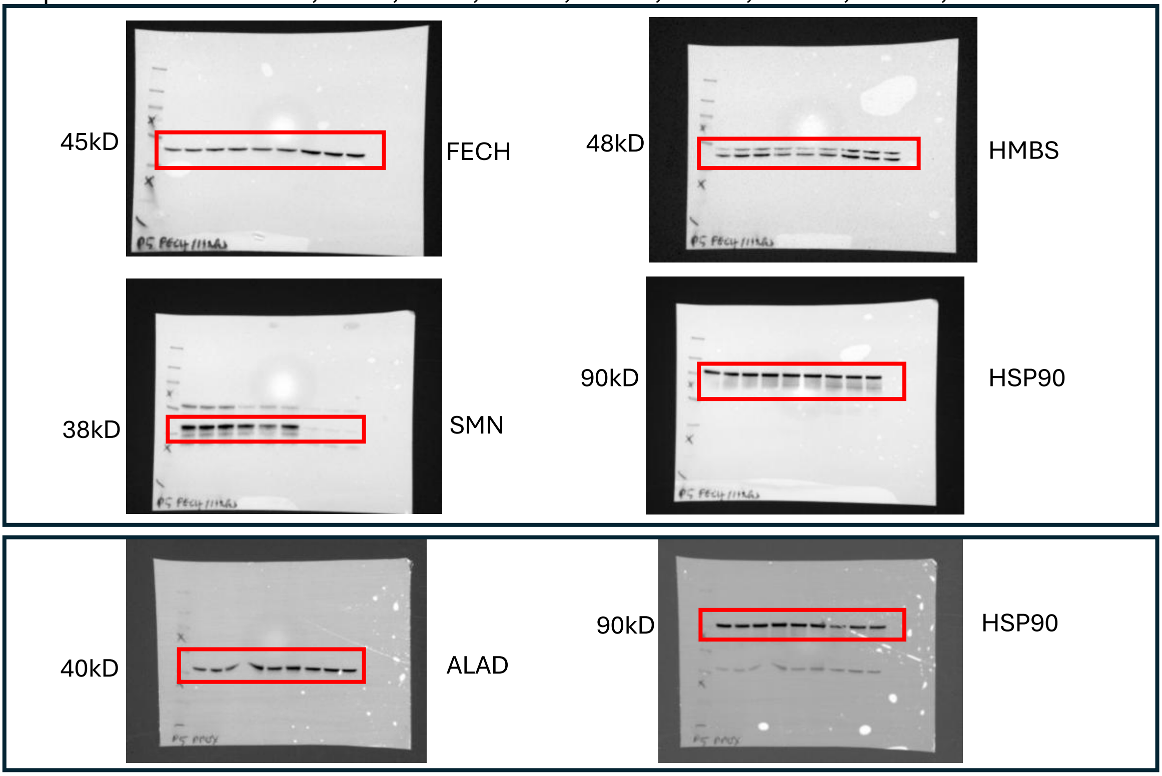

#### Figure 5A

Samples lane order: WT-1, WT-2, WT-3, HET-1, HET-2, HET-3, SMA-1, SMA-2, SMA-3, SMA-4.

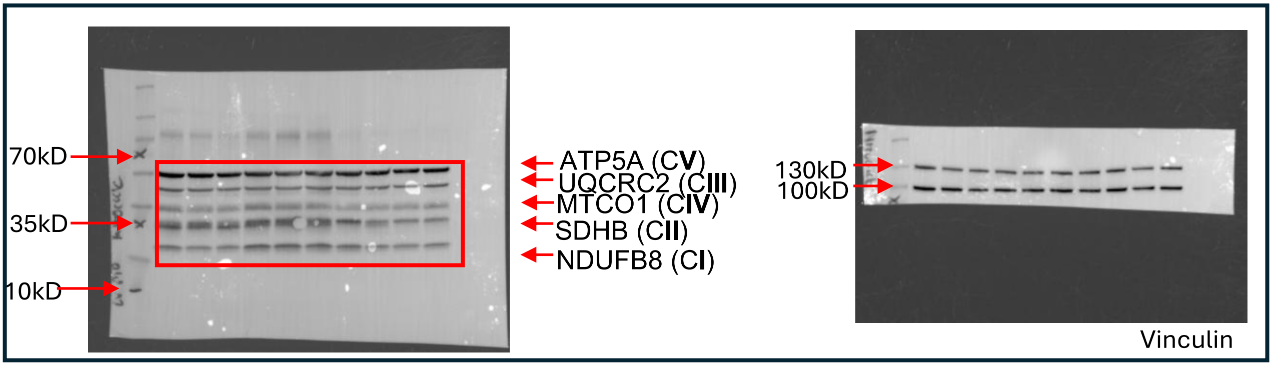


#### Figure 5B

Samples lane order: Whole liver homogenate 1, 2, 3, isolated mitochondria 1, 2, 3.

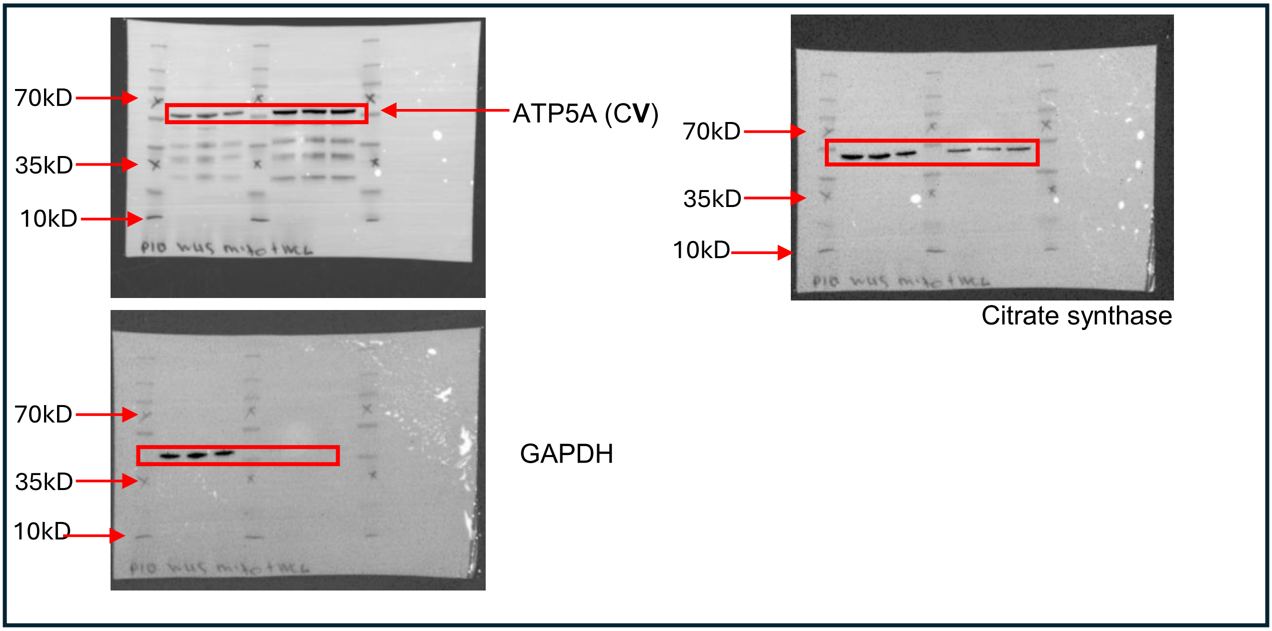


#### Figure 5G

Samples lane order: WT-1, WT-2, WT-3, HET-1, HET-2, HET-3, SMA-1, SMA-2, SMA-3, SMA-4.

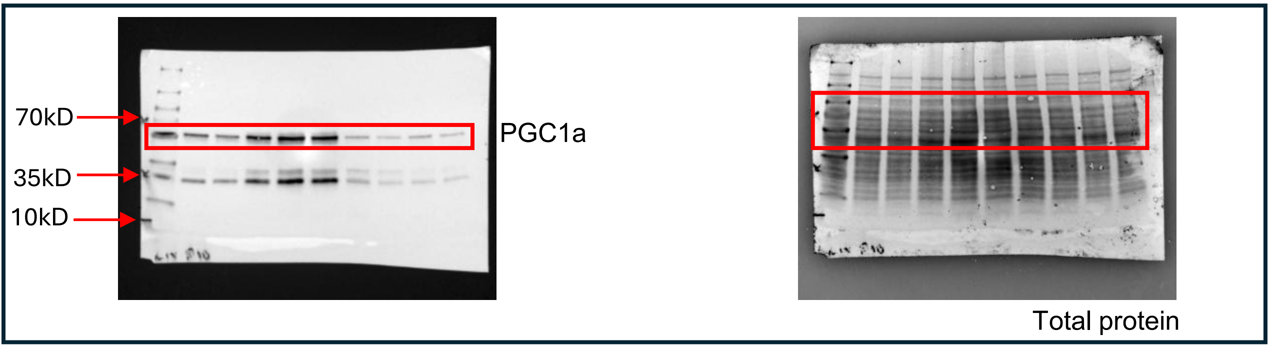

### Figure 5D

Samples lane order: WT, HET, SMA, SMA+ASO

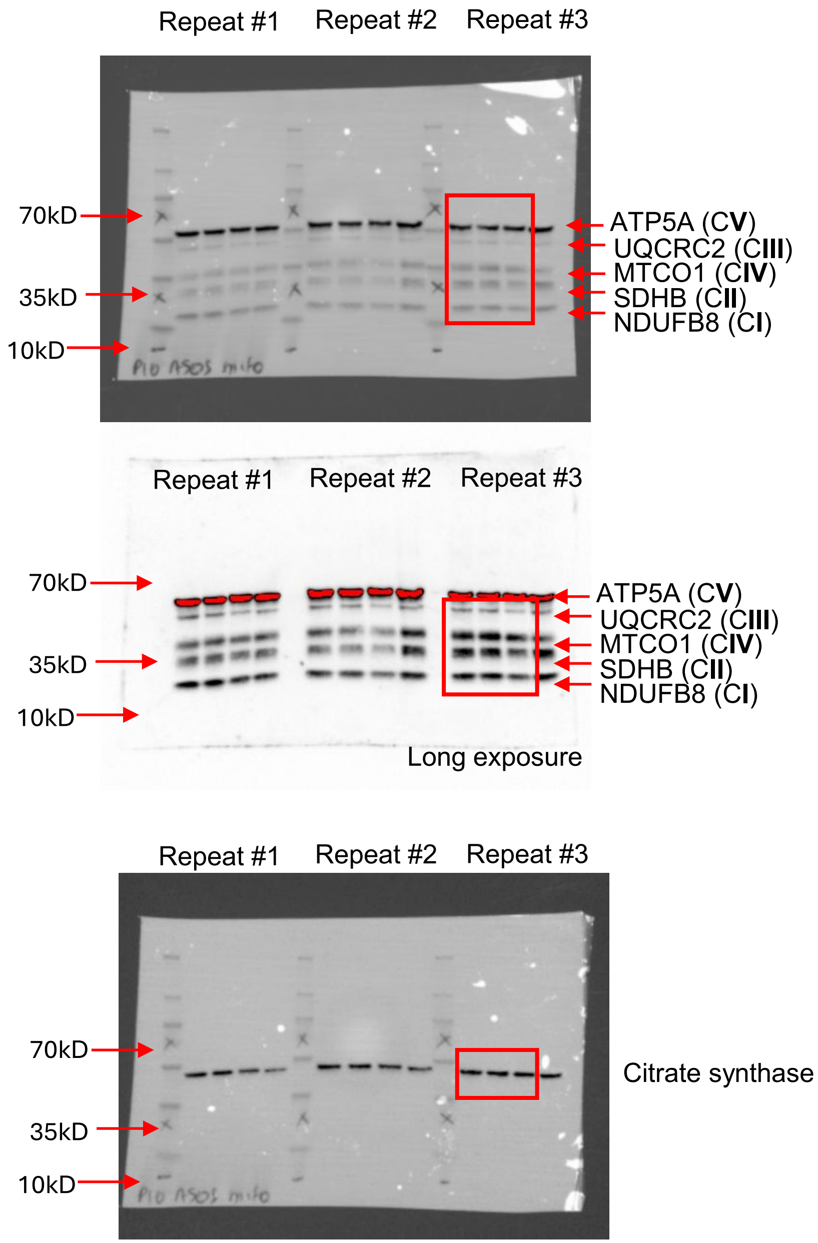

### Figure 6A

Samples lane order: WT-1, WT-2, WT-3, HET-1, HET-2, HET-3, SMA-1, SMA-2, SMA-3, SMA-4.

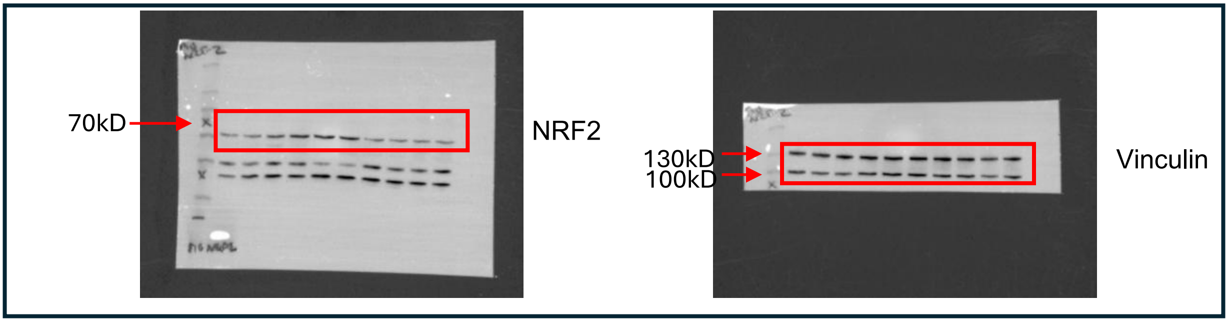


### Figure 6A

Samples lane order: WT-1, WT-2, WT-3, HET-1, HET-2, HET-3, SMA-1, SMA-2, SMA-3.

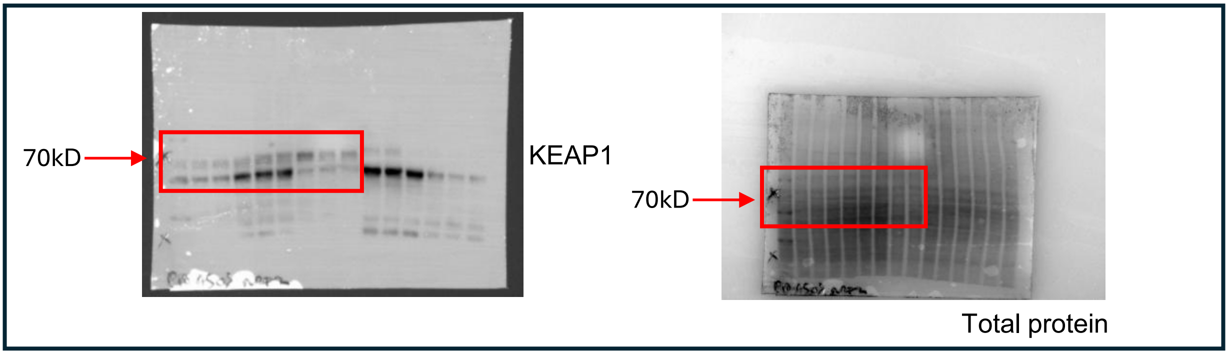


### Figure 6D

Samples lane order: WT-1, WT-2, WT-3, HET-1, HET-2, HET-3, SMA-1, SMA-2, SMA-3, SMA-4.

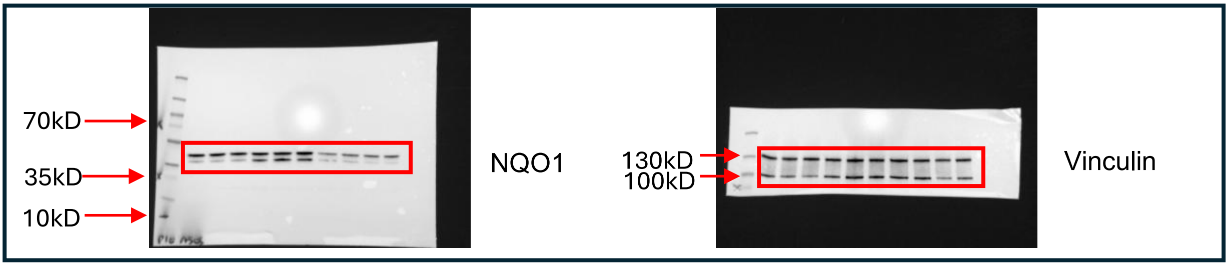


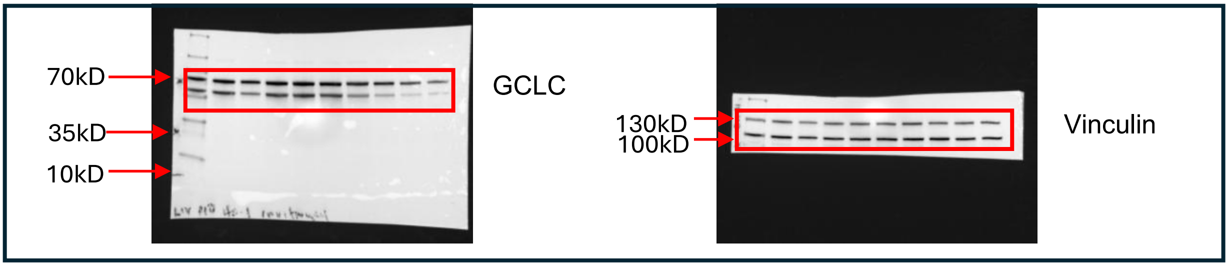


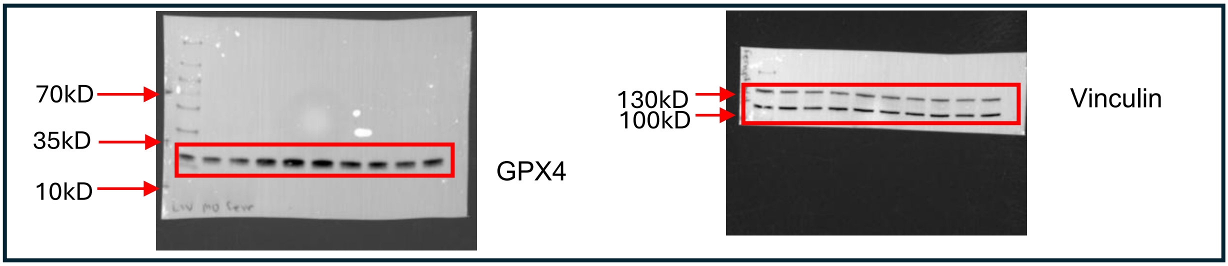

#### Figure 6I

Samples lane order: WT-1, WT-2, WT-3, HET-1, HET-2, HET-3, SMA-1, SMA-2, SMA-3, SMA-4.

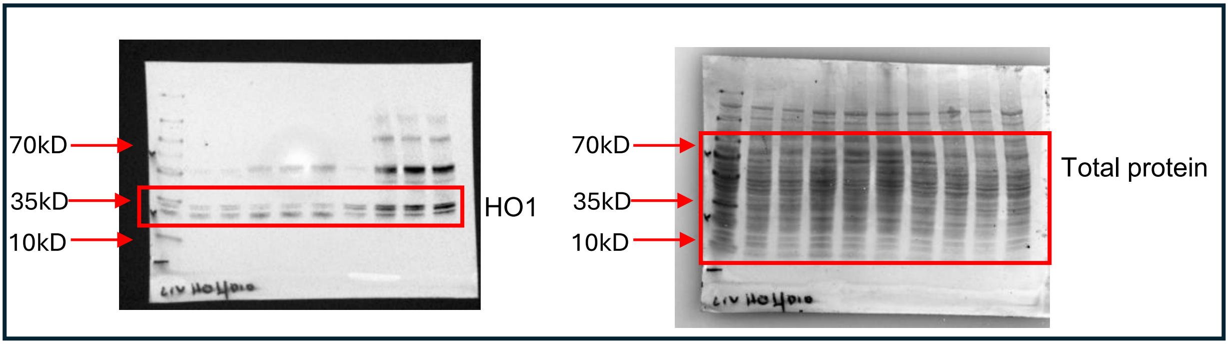


#### Figure 7B

Samples lane order: WT-1, WT-2, WT-3, HET-1, HET-2, HET-3, SMA-1, SMA-2, SMA-3, HET+ASO-1, HET+ASO-2, HET+ASO-3, SMA+ASO-1, SMA+ASO-2, SMA+ASO-3.

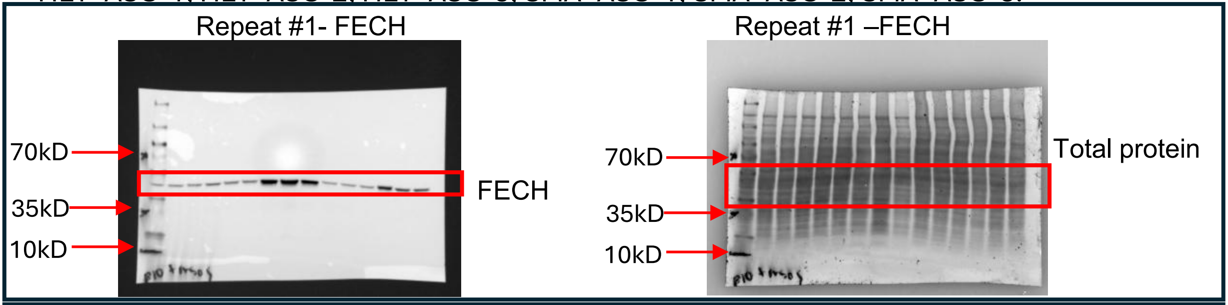


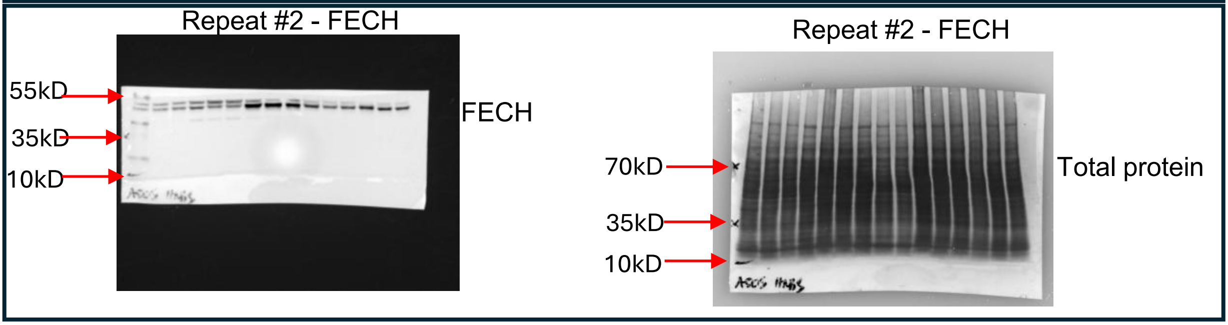


#### Figure 7B

Samples lane order: WT-1, WT-2, WT-3, HET-1, HET-2, HET-3, SMA-1, SMA-2, SMA-3, HET+ASO-1, HET+ASO-2, HET+ASO-3, SMA+ASO-1, SMA+ASO-2, SMA+ASO-3.

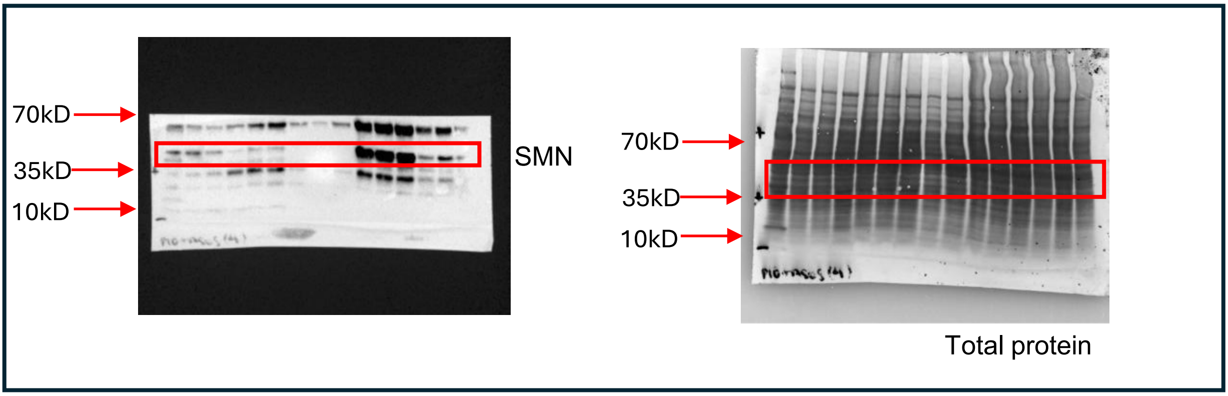

### Figure 8B, C

Samples lane order: WT, HET, SMA, SMA+ASO

CBB-BNPAGE → OXPHOS-BNPAGE → CI- CNPAGE → CIV – CN-PAGE → CII – CN- PAGE

Repeat #1-CBB

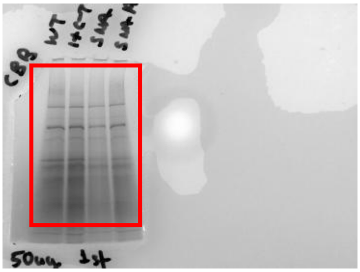

Repeat #1-OXPHOS

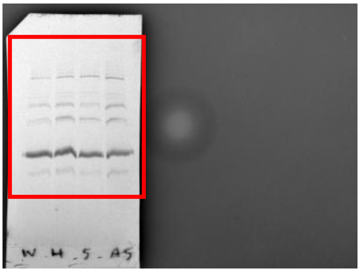

Repeat #1-CI

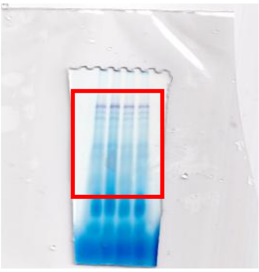

Repeat #1-CIV

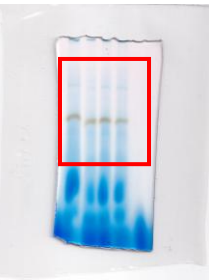

Repeat #2-CBB

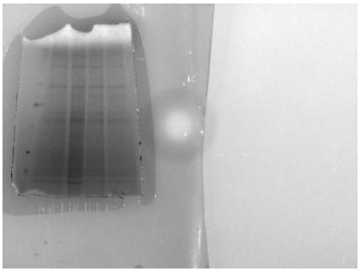

Repeat #2-OXPHOS

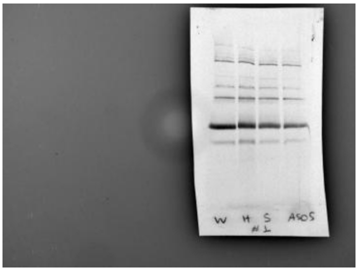

Repeat #2-CI

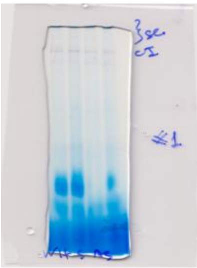

Repeat #2-CIV

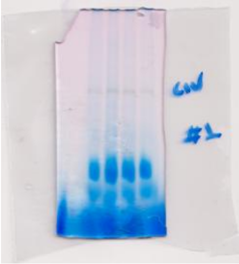

Repeat #3-CBB

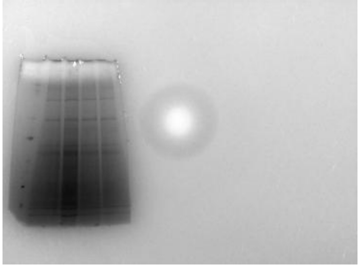

Repeat #3-OXPHOS

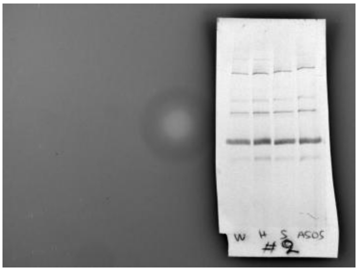

Repeat #3-CI

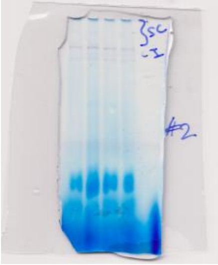

Repeat #3-CIV

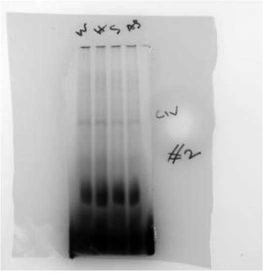

Repeat #3-CI

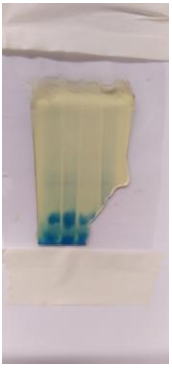

Repeat #3-CI

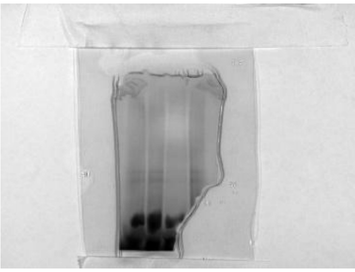

Repeat #1, #2 -CI

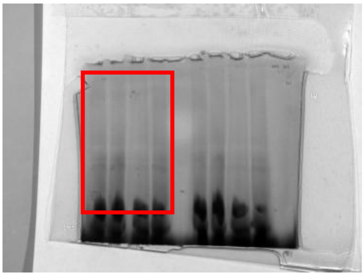

Repeat #1, #2 -CI

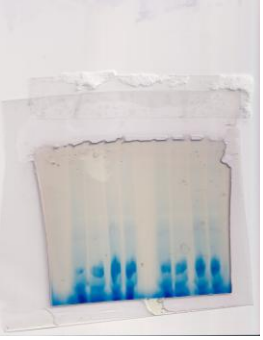

#### Figure 8H

Samples lane order: WT-1, WT-2, WT-3, HET-1, HET-2, HET-3, SMA-1, SMA-2, SMA-3, HET+ASO-1, HET+ASO-2, HET+ASO-3, SMA+ASO-1, SMA+ASO-2, SMA+ASO-3.

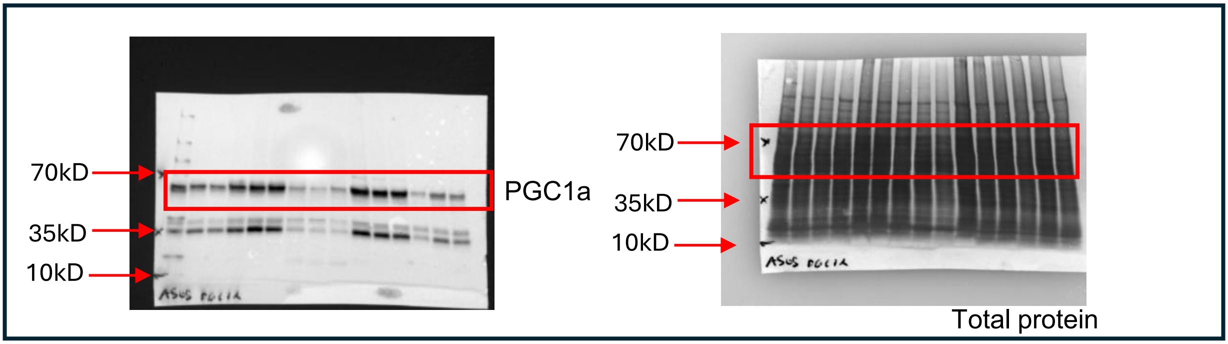


#### Figure 9A

Samples lane order: WT-1, WT-2, WT-3, HET-1, HET-2, HET-3, SMA-1, SMA-2, SMA-3, HET+ASO-1, HET+ASO-2, HET+ASO-3, SMA+ASO-1, SMA+ASO-2, SMA+ASO-3.

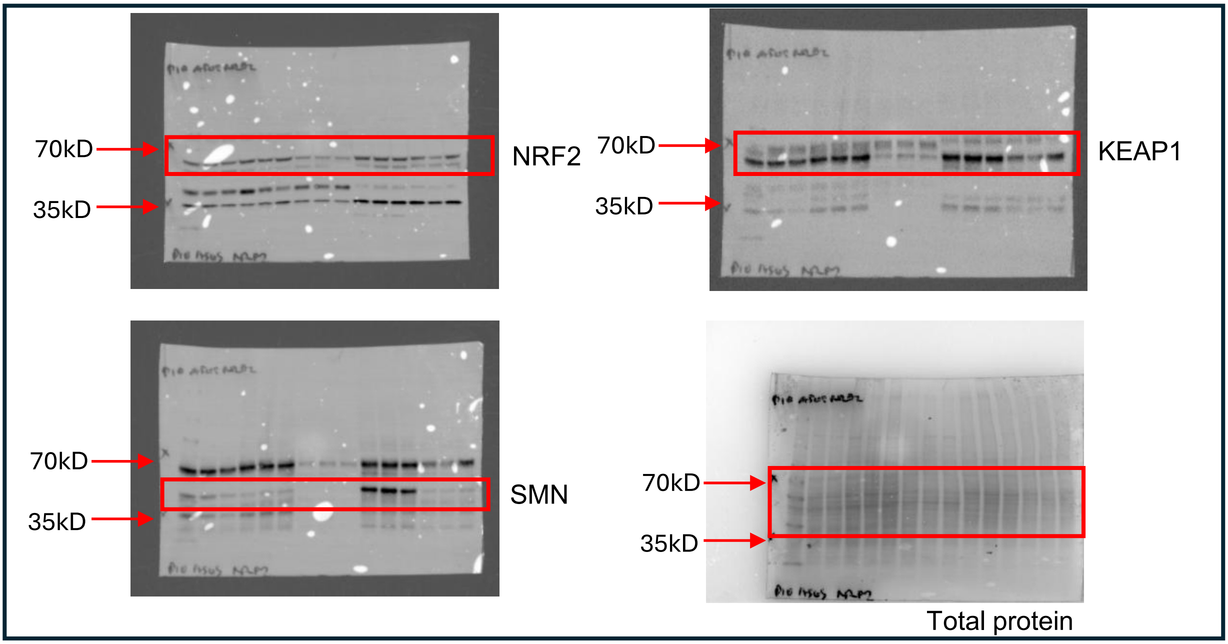

#### Figure 9D

Samples lane order: WT-1, WT-2, WT-3, HET-1, HET-2, HET-3, SMA-1, SMA-2, SMA-3, HET+ASO-1, HET+ASO-2, HET+ASO-3, SMA+ASO-1, SMA+ASO-2, SMA+ASO-3.

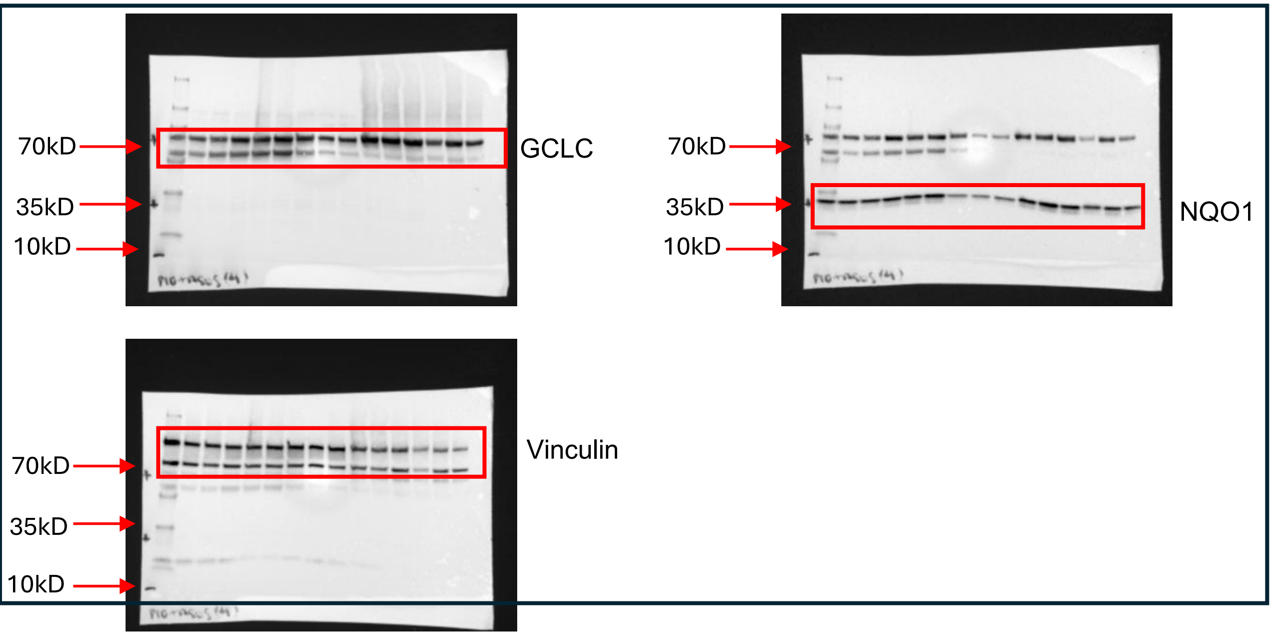


#### Figure 9G

Samples lane order: WT-1, WT-2, WT-3, HET-1, HET-2, HET-3, SMA-1, SMA-2, SMA-3, HET+ASO-1, HET+ASO-2, HET+ASO-3, SMA+ASO-1, SMA+ASO-2, SMA+ASO-3.

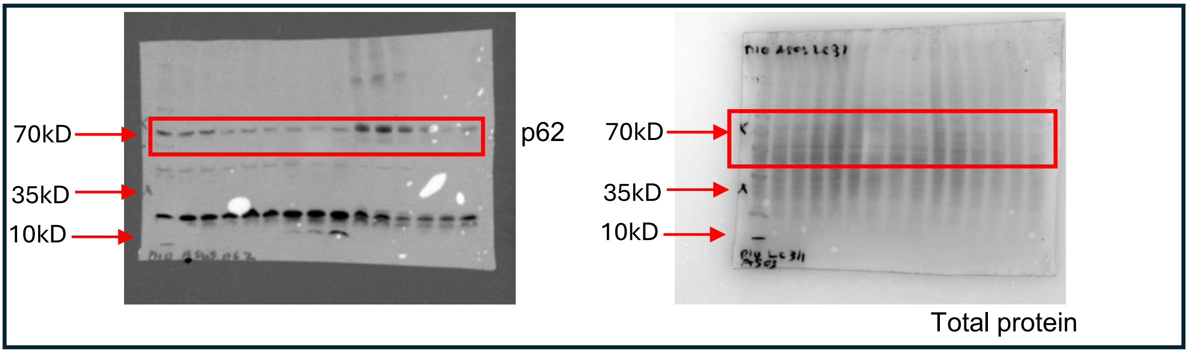

#### Supplementary Figure S8A.

Samples lane order: WT-1, WT-2, WT-3, HET-1, HET-2, HET-3, SMA-1, SMA-2, SMA-3.

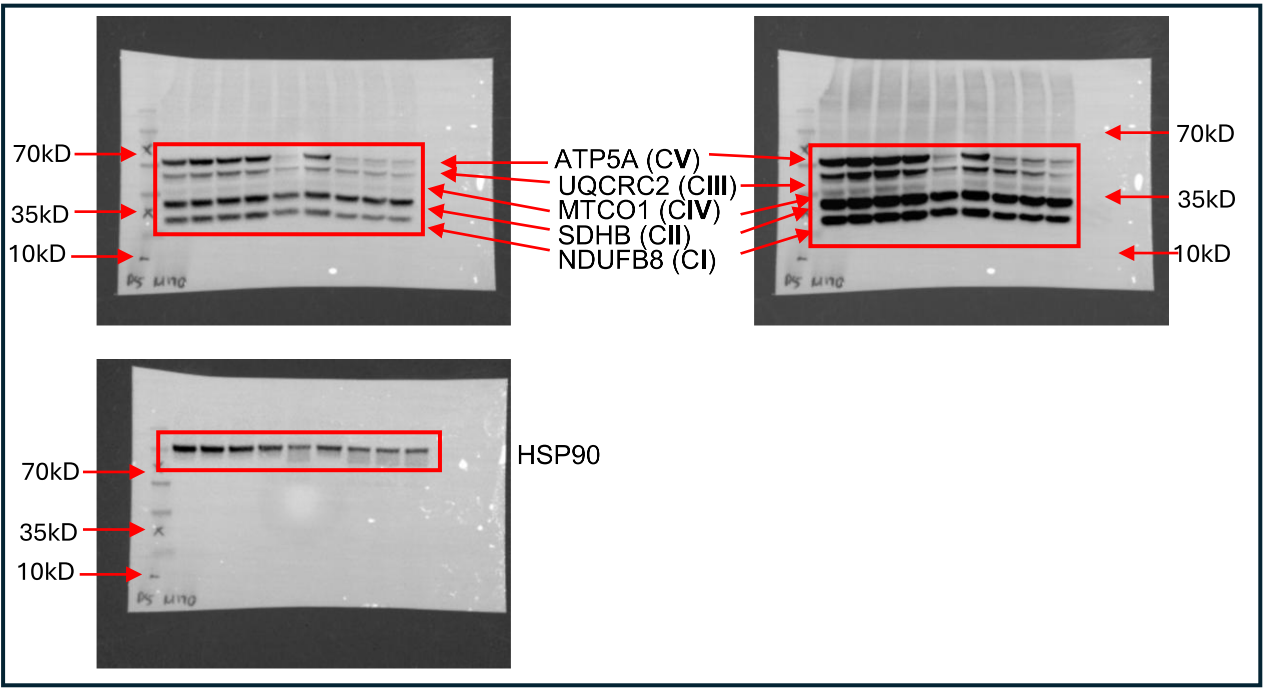


#### Supplementary Figure S8B.

Samples lane order: WT-1, WT-2, WT-3, HET-1, HET-2, HET-3, SMA-1, SMA-2, SMA-3.

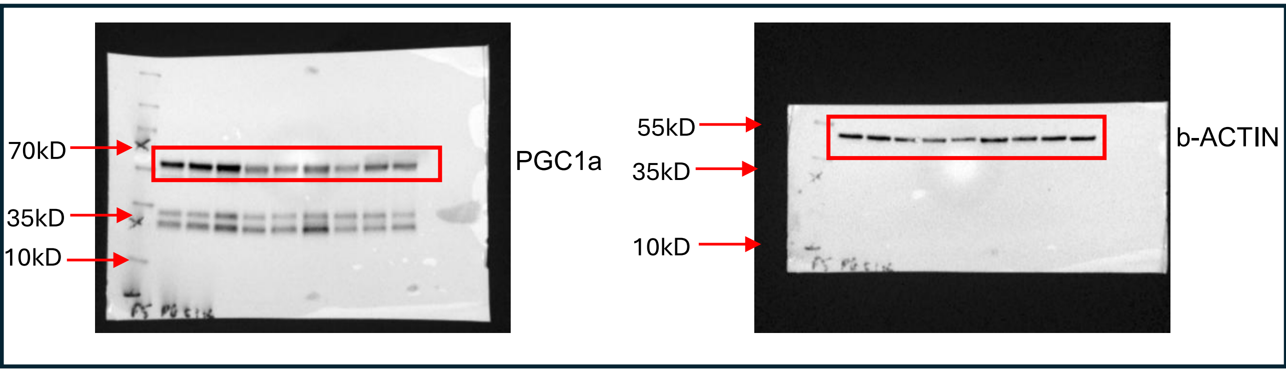

### Supplementary Figure S10A.

Samples lane order: WT-1, WT-2, WT-3, HET-1, HET-2, HET-3, SMA-1, SMA-2, SMA-3.

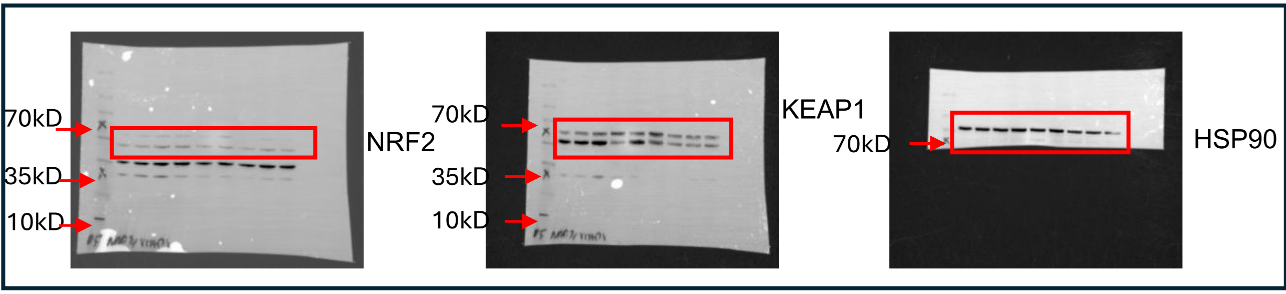


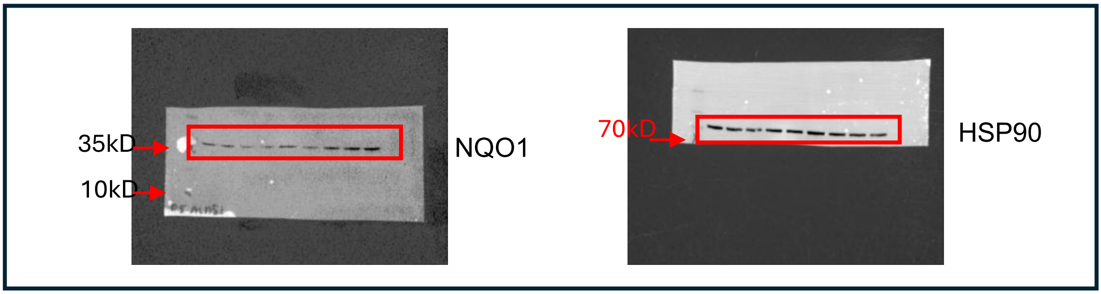


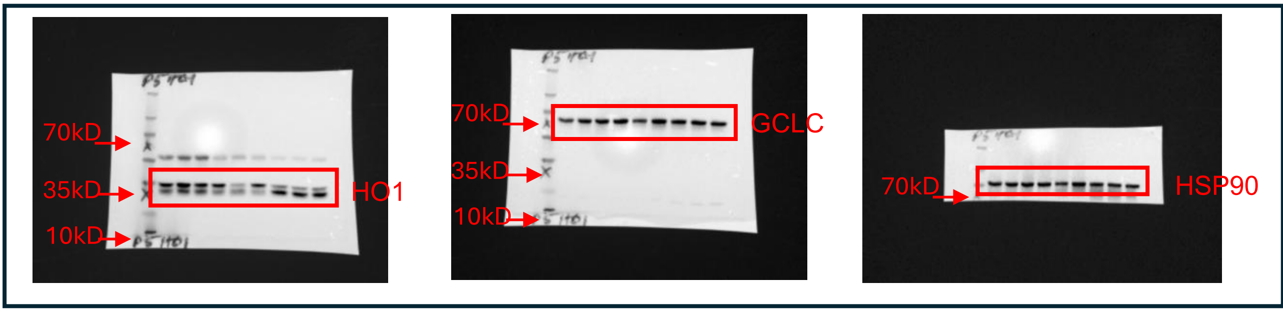


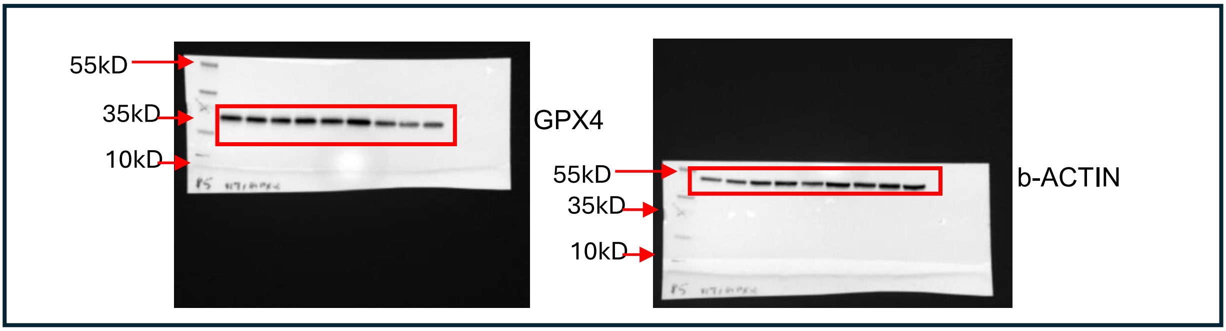

### Supplementary Figure S11A

Samples lane order: HET-1. HET-2, HET-3, SMA-1, SMA-2, SMA-3, HET+ASO-3, SMA+ASO-1, SMA+ASO-2, SMA+ASO-3.

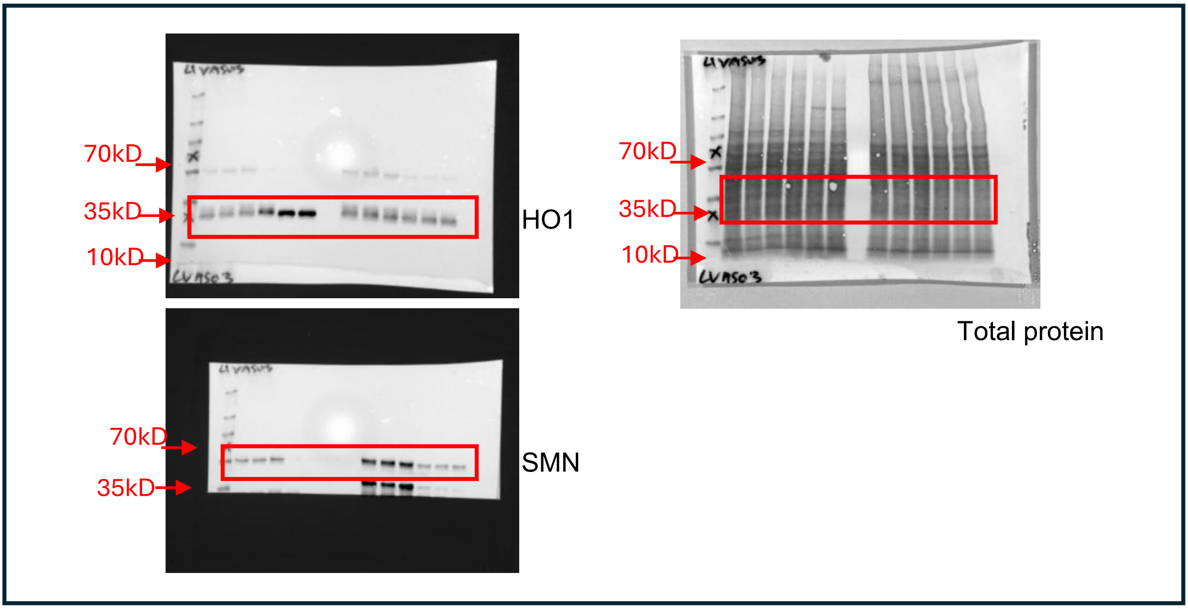


### Supplementary Figure S12A

Samples lane order: WT-1, WT-2, WT-3, HET-1. HET-2, HET-3, SMA-1, SMA-2, SMA-3, HET+ASO-3, SMA+ASO-1, SMA+ASO-2, SMA+ASO-3.

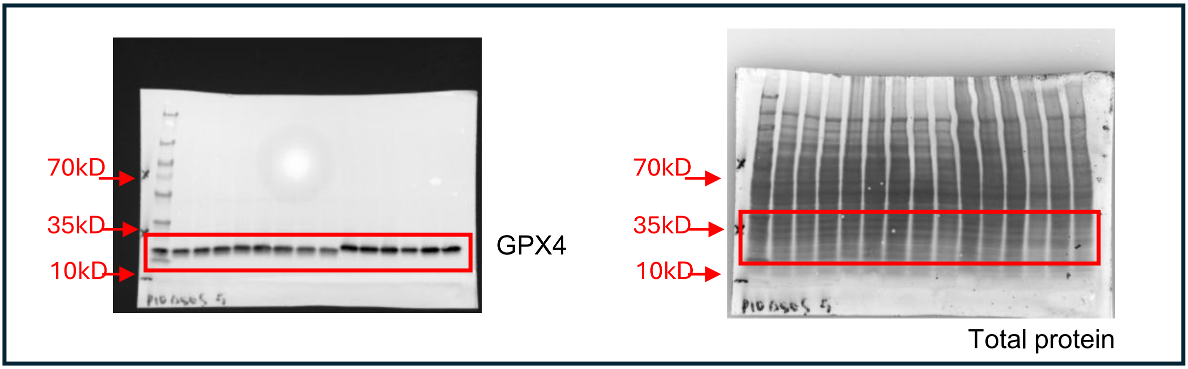
